# Longitudinal flux balance analyses of a patient with Crohn’s disease highlight microbiome metabolic alterations

**DOI:** 10.1101/2022.12.19.520975

**Authors:** Arianna Basile, Almut Heinken, Johannes Hertel, Larry Smarr, Weizhong Li, Laura Treu, Giorgio Valle, Stefano Campanaro, Ines Thiele

## Abstract

Inflammatory bowel diseases (IBD) are characterised by episodic inflammation of the gastrointestinal tract. Gut microbial dysbiosis characterises the pathoetiology, but its role remains understudied. We report the first use of constraint-based microbial community modelling on a single individual with IBD, covering seven dates over 16 months, enabling us to identify a number of time-correlated microbial species and metabolites. We find that the individual’s dynamical microbial ecology in the disease state drives time-varying *in silico* overproduction, compared to healthy controls, of more than 24 biologically important metabolites, including oxygen, methane, thiamine, formaldehyde, trimethylamine N-oxide, folic acid, serotonin, histamine, and tryptamine. A number of these metabolites may yield new biomarkers of disease progression. The microbe-metabolite contribution analysis revealed that some genus *Dialister* species changed metabolic pathways according to the disease phases. At the first time point, characterised by the highest levels of blood and faecal inflammation biomarkers, they produced L-serine or formate. The production of the compounds, through a cascade effect, was mediated by the interaction with pathogenic *Escherichia coli* strains and *Desulfovibrio piger*. We integrated the microbial community metabolic models of each time point with a male whole-body, organ-resolved model of human metabolism to track the metabolic consequences of dysbiosis at different body sites. The presence of *D. piger* in the gut microbiome influenced the sulphur metabolism with a domino effect affecting the liver. These results underline the importance of tracking an individual’s gut microbiome along a time course, creating a new analysis framework for self-quantified medicine.

**Graphical abstract:** 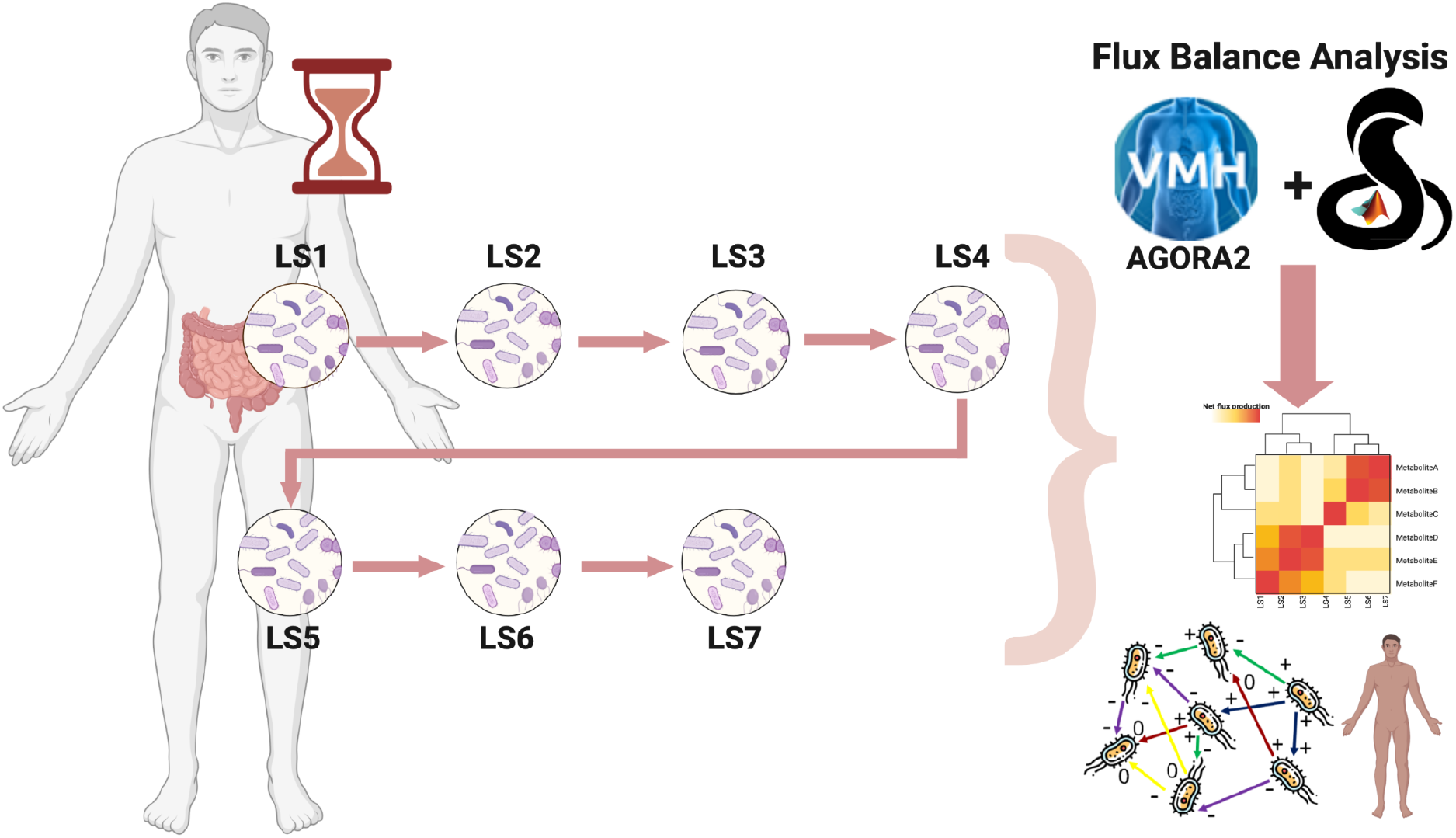

## Introduction

The human gut microbiome performs essential functions in shaping the host immune system, host cell proliferation, and is involved in the maintenance of endocrine functions ^1^. The human microbiome consists of a large number of archaeal and bacterial, but also of viral and fungal, species ^2^. The composition of the microbiome depends on host factors, such as age, sex, location, ethnicity, and lifestyle (e.g., diet, exercise, and medication). Between healthy individuals, the relative abundances of taxa are highly variable, while the functional capabilities are more stable. In contrast, many multifactorial diseases are characterised by a dysbiotic microbiome ^3^.

The gut microbiota differs among individuals, has a variable composition in different parts of the digestive tract, and can undergo extensive modifications throughout life ^4^. This aspect is regarded as an obstacle to gut microbiome-based medical applications, as it remains difficult to identify a clear signature of the dysbiotic microbiota. The microbiome of an individual can change during the outbreak of a disease, and the symptomatology of the patient changes accordingly ^5^. This is particularly true for patients affected by Inflammatory Bowel Disease (IBD), a disorder characterised by episodic symptoms driven by time-changing inflammation of the gastrointestinal tract ^6^. This time-varying inflammation is the byproduct of the constant interaction between the human host’s immune system and the changing ecological profile of the host’s gut microbiome ^7^. The presence or absence of inflammation is strongly associated with four measurable faecal biomarkers: calprotectin and lactoferrin (shed from white blood cells), lysozyme (innate immune system), and secretory IgA (adaptive immune system) ^8^, as indicators of levels of severity of episodic IBD. Historically, IBD has been considered to have two main subtypes: ulcerative colitis (UC) and Crohn’s disease (CD). However, a large (30,000 patients) human genotype study in 2016 ^9^ demonstrated that the human genetic predisposition is best explained by three subtypes: ileal Crohn’s disease (ICD), colonic Crohn’s disease (CCD), and UC. This same IBD tripartite division is seen when the gut microbiome ecology is clustered ^10^. This separation into three subtypes is even clearer when using the Kyoto Encyclopaedia of Genes and Genomes (KEGG) database to cluster the gut microbiome of patients ^11^.

Recently, the study of the microbiome has moved from “Who is there?” to “What are they doing?”. In particular, the constraint-based reconstruction and analysis (COBRA) framework, which relies on a genome-scale reconstruction of a target organism’s metabolism and the application of condition-specific constraints, e.g., meta-omics data and allowed uptake of nutrients ^12^, has moved these questions further to “What do they produce?”, and “How do they interact?”^1,13–16^. Genome-scale reconstructions are assembled using organisms’ genome sequences and biochemical, genetic, and physiological evidence ^17^. COBRA assumes the biological systems to be at a steady state, i.e., the change in metabolite concentration over time is zero. Flux balance analysis (FBA) ^16^, a frequently used COBRA method, assumes in addition that the biological system tries to achieve an objective, e.g., maximal biomass yield ^18^. FBA has been successfully applied to investigate the role of the human gut microbiome in various complex diseases, including Parkinson’s disease ^19,20^, and inflammatory bowel disease ^1,21–23^. To facilitate the application of constraint-based modelling to research on the human gut microbiome, the AGORA (Assembly of Gut Organisms through Reconstruction and Analysis) collection was established ^24^, and recently expanded to cover over 7,200 semi-manually curated microbial genome-scale metabolic reconstructions ^25^.

In prior studies, FBA was used on a set of microbiome samples comparing healthy individuals with IBD patients at a single time per patient. COBRA modelling has been used to link mechanistically host-microbiome-environment interactions to IBD-related changes ^12^. The potential of 818 microbial strains to deconjugate primary bile acids into secondary bile acids has been investigated with a combined approach based on comparative genomics followed by FBA ^26^. In that study, it has been reported that microbial species can complement each other’s bile acid pathway to achieve the broader bile acid production repertoire observed in faecal samples ^26^.

Despite the numerous studies performed on CD, the evolution of the gut microbiome during disease onset and progression has not been analysed with FBA. Here, we bridge this gap by investigating how the extreme dysbiosis time variations of the gut microbiome ecology in a single individual with CCD can cause similar large time variations in a number of key metabolites. Another unexplored aspect of the disease is the interaction between host metabolism and the compounds produced by the normal and the dysbiotic gut microbiome. To tackle this specific aspect, we performed an additional investigation using sex-specific, organ-resolved, whole-body metabolic models of human metabolism, which account for 28 organs, tissues, and cell types ^27^.

The present study used FBA to analyse the metabolic evolution of the gut microbiome community in a single individual (“LS”) affected by left-sided CCD across seven time points covering a period of 16 months in 2012/2013 (Fig. 1). The metagenomic data for these seven time points have been previously compared ^11,28^ with a set of metagenomic data from healthy individuals drawn from the NIH Human Microbiome Project ^29^, as well as selected metagenomic data from patients with ICD and with UC. It has been shown ^30^ that LS’s gut microbiome taxonomic profile deviated a great deal from the healthy individuals and furthermore, that his gut microbiome exhibited major taxonomic shifts over time.

**Figure 1:**
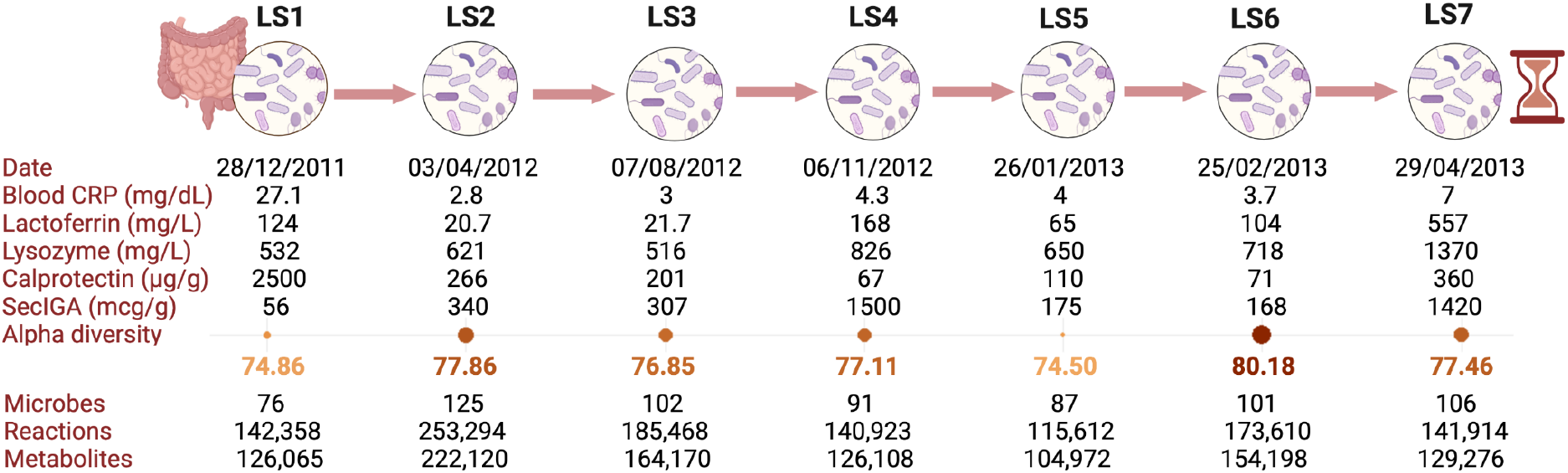
Timeline with metadata of the different samples. In the timeline, generated with BioRender, the collection date, the measured blood concentration of complex reactive protein (CRP), as well as faecal lactoferrin, lysozyme, calprotectin, and secretory IgA (SecIGA) are reported. LS has episodic major increases in all of these inflammatory/immune biomarkers, as healthy values for each are CRP<1, lactoferrin<7.3, lysozyme<600, calprotectin<50, and SecIGA (30-275). The medical intervention between LS1 and LS2 consisted of ciprofloxacin, metronidazole, and prednisone. “Microbes” refers to the number of metabolic models identified in the metagenomic samples according to the threshold selected (see Methods) and that were included in each time point-specific microbial community model. The number of reactions and metabolites refers to the size of the condition-specific microbial community models at each time point.

In the present study, the metagenomic data for LS’s seven time points, computed at both the species and strain levels (Supplementary Table S1) were mapped onto the AGORA2 collection ^25^. The taxonomic composition of the microbiome samples in AGORA2 for each time point (Supplementary Table S1) was then analysed and diversity indexes were computed. As a comparison, we used the metagenomic relative abundance of both species and strain data from the cohort of 34 healthy control subjects (Supplementary Table S1) and also mapped them on AGORA2 (Supplementary Table S1). Subsequently, the contribution of each microbial species within the microbiome-level, time point-specific AGORA2 microbial community models to the global profile of 740 metabolites (Supplementary Table S2) present in the gut was predicted (Methods: Simulations section).

We found that LS’s major gut microbiome taxonomic shifts over time led to correspondingly large FBA metabolic shifts from the personalised microbial community models. In particular, our model results show that a number of biologically important metabolites were highly (10-10,000x) overproduced, compared to HeAve, at various time points in LS’s samples, including oxygen, methane, thiamine, formaldehyde, trimethylamine N-oxide, folic acid, serotonin, histamine, and tryptamine. Furthermore, our results suggest that through the production of few metabolites, i.e., L-serine and formate, species of the *Dialister* genus cooperate with many pathogenic strains, such as adherent invasive *Escherichia coli* strains, archaeal species, and *Desulfovibrio piger* ATCC2. The interactions trigger inflammatory responses and enhance methane production. Finally, *D. piger* ATCC2 plays an important role in the production of the host-toxic SO_3_^2-^. Additionally, we investigated host-microbiome co-metabolism during these time points. In conclusion, our study sheds light on metabolites and microbial species triggering the inflammatory responses, and their impact on host metabolism.

## Results and discussion

### Characterisation of the time points

The n=1 patient “LS” with CCD was a non-smoker male, 64 years old at baseline. A detailed description of his medical history is reported in Supplementary Materials. We carried out (see Methods) deep (~100M reads per sample) shotgun metagenomic sequencing, yielding 510 species and 790 strains relative abundance, on frozen faecal samples for seven time points, deemed LS1 to LS7 (Supplementary Table S1) according to the time of collection.

All time points were characterised by abnormal concentrations of hematic and faecal immune or inflammatory biomarkers, with LS1 having both the highest hematic complex reactive protein (CRP) and the highest faecal calprotectin concentration (**Fig. 1**, Supplementary Table S1). In contrast, lactoferrin, lysozyme, and secretory IGA had their highest values at LS4 and LS7. The medical intervention between LS1 and LS2 (**Fig. 1**) consisted of two antibiotics, being 500 mg ciprofloxacin administered orally twice a day and 250 mg metronidazole administered orally three times per day for one month starting from 31^st^ of January 2012 ^31^. During this period, the patient also received daily 40 mg oral prednisone, a drug used to suppress the immune system and decrease inflammation.

In addition, we obtained shotgun metagenomic data (^~^100M reads per person) for the gut microbiome from 34 healthy individuals in the Human Microbiome Project ^32^ (Methods, Supplementary Table S1). This control dataset allowed for the comparison of the LS microbiome with healthy individuals and for the identification of microbial and functional differences associated with the disease status at each time point.

### Analysis of metagenomic data with microbiome-level metabolic models

First, we investigated the changes in the metagenomic phyla abundances in the healthy and disease microbiomes. We identified major differences over time between the seven LS times and the healthy microbiomes across the seven most abundant phyla: Actinobacteria, Bacteroidetes, Euryarchaeota, Firmicutes, Fusobacteria, Proteobacteria and Verrucomicrobia (**Fig. 2A**, Supplementary Table S1). Then, we used the strains identified in the shotgun metagenomic data of LS (Supplementary Table S1) as input to the AGORA2 collection of microbial metabolic reconstructions ^25^ (see Methods). This process creates seven *in silico* microbial community models (Supplementary Table S1) accounting for a total of 214 distinct microbes, covering both bacterial and archaeal species. The simulation (Methods) uses the seven ecological models to compute the metabolites produced by the microbial communities. In detail, the maximal production and uptake fluxes of each metabolite from all the microbial species is computed, following ^16^. In the following sections, we will state when we are referring to the input metagenomic microbial abundances or to the AGORA2 mapped abundances.

**Figure 2:**
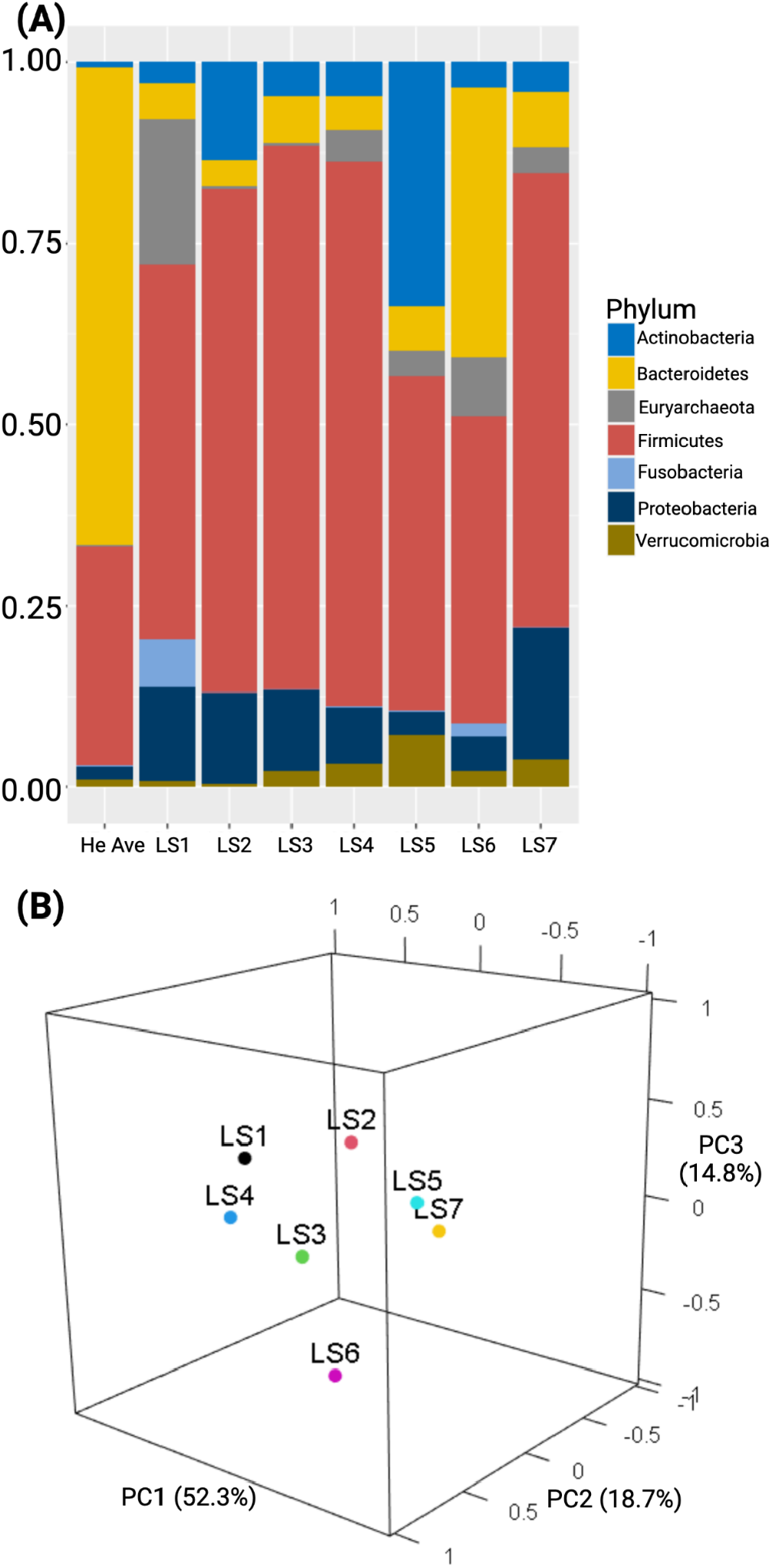
Graphical representation of the time evolution of the gut microbiome ecology. **(A)**Stacked barplot representing the metagenomic phyla abundance of the gut microbiome in the different LS samples with a comparison to the Healthy Average (HeAve). For a similar barplot, which also shows the time variation of 10 abundant species superimposed on the LSphyla bars, see Fig. 2 in Ref 30. **(B)**3D principal component analysis computed on the species abundances mapped onto AGORA2 involved in the metabolic modelling for each time step and reflecting differential microbial compositions and abundances is shown (more details in the Method section). For a 2D PCA of the metagenomic species relative abundances of the 7 LS samples and the 34 HE samples, see Figure 4a of Ref. 11.

Using the metagenomic relative abundances, we compared LS microbial species composition at each time point and with those of the 34 healthy controls, including calculating both the maximum (HeMax) and average (HeAve) abundance of each microbial species across the healthy individuals (Supplementary Table S1). We observed cases where LSMax>HeAve, meaning that the disease state associated microbiome fluctuated over time and exceeded the average relative abundance in the healthy population. Additionally, we identified cases with LSMax>HeMax, meaning that the relative abundance in the disease state could be greater than the largest cross-population variation. Using this comparison, the dysbiosis experienced by LS was characterised by a major decrease in microbe species that were dominant in the healthy individuals, thereby allowing for the time-dependent bloom of typically less abundant microbes in LS1-7 (Supplementary Fig. S3-S9).

In more detail, the average healthy control’s gut microbiomes were found to be predominantly composed of Bacteroidetes (65.6%) and a lower fraction of Firmicutes (30%) (**Fig. 2A**). In contrast, the most abundant LS microbiome phylum at all seven time points was Firmicutes, which ranged from 1.4 to 2.5x the HeAve abundance (**Fig. 2A,** Supplementary Table S1). The overabundance of Firmicutes was driven by the blooming of normally rare Firmicutes species from classes Bacilli and Clostridia, with overabundances ranging from 100-1,000x HeAve for those species. In particular, the family Lachnospiraceae (in class Clostridia) was 2.6-3.7x HeAve, mainly represented by *Dorea longicatena* DSM 13814, normally rare, but 25x HeAve in LS4 (**Fig. 3**, Supplementary Table S1). In contrast, the other dominant microbial phylum in healthy individuals, Bacteroidetes, was depleted by more than 10x in all LS time points except LS6 when it bounced back to half the abundance of HeAve.

**Figure 3:**
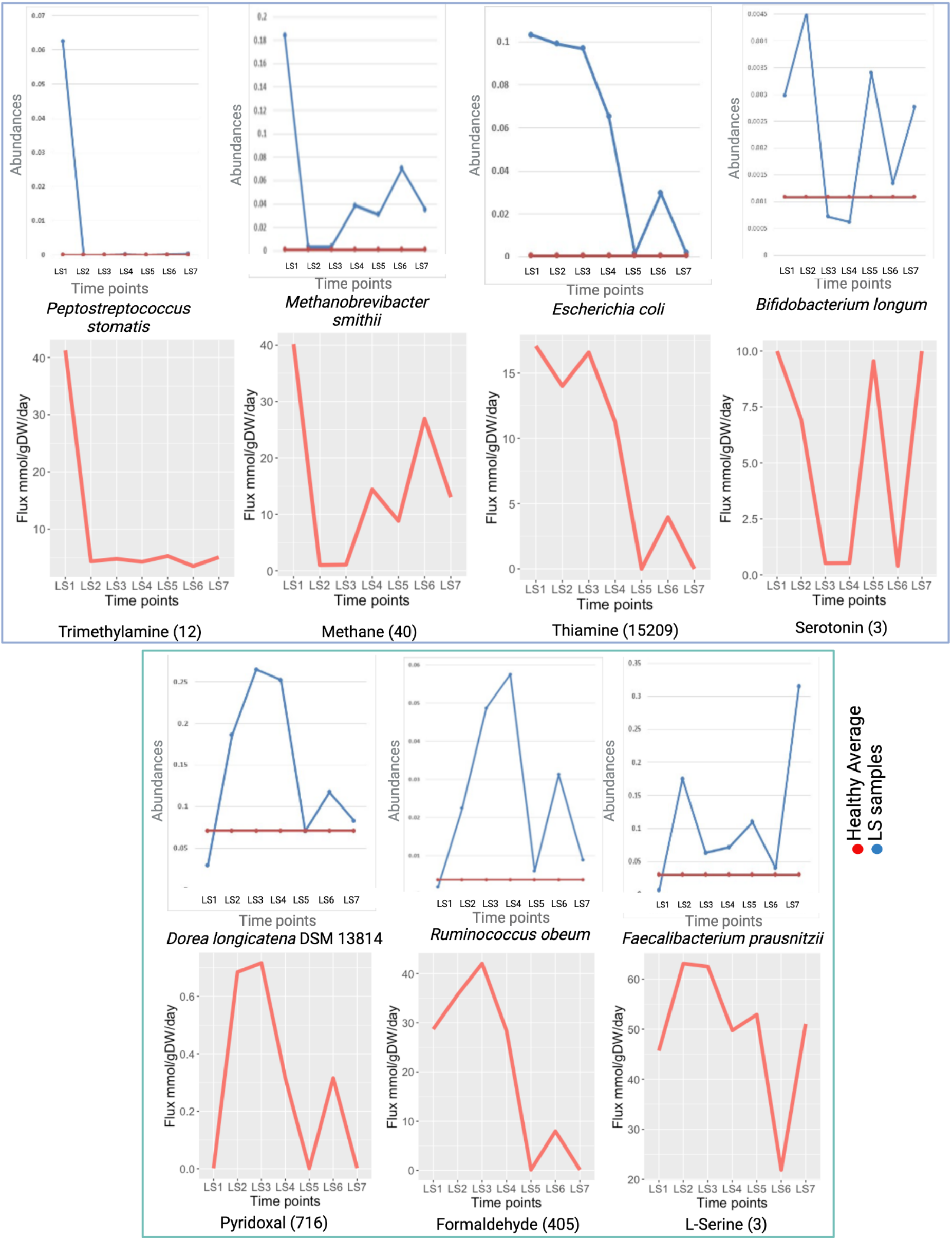
Correlations between microbial abundances and secretion fluxes of key metabolites. **Top square**: Class I microbe-metabolite relationships. **Bottom square**: Class II. Each pair of graphs represents representative specific gut microbiome species relative abundance over LS1-7 (top of pair) and a matched metabolite flux over LS1-7 (bottom of pair). For the species graph, the blue line represents the relative abundance of the microbe over LS1-7, while the red line represents the relative abundance of that microbe for HEAve.

The ecological absence of the normally dominant phylum Bacteroidetes allowed other, normally rare phyla in the healthy individuals, to dynamically bloom. In particular, the phylum Euryarchaeota was elevated by at least three times in all samples when compared with the HeAve, with an extreme overabundance in LS1 and LS6, which are 137x HeAve and 57x HeAve, respectively (Supplementary Table S1). The observed high archaeal relative abundances in all the phases are typical of CD-associated dysbiosis ^33^. In particular, the presence of the family Methanobacteriaceae (dominated by *Methanobrevibacter smithii*) was strongly influenced by the disease, varying between 3-170x HeAve (Supplementary Fig. S13), with the highest value occurring at LS1. The phylum Proteobacteria was also overabundant, compared to healthy individuals, at all seven time points. For LS1-3, it was approximately seven times higher, and for LS7, it was ten times higher than HeAve. Within this phylum, the family Enterobacteriaceae reached a peak of >150x HeAve in LS7 (Supplementary Table S1, Supplementary Fig. S10). The abundance of family Enterobacteriaceae species *E. coli* in LS1 was 187x (Supplementary Table S1, Supplementary Fig. S11). Phylum Actinobacteria had a higher abundance (4-50x HeAve) at all time points and a higher diversity with 58 different species present at time point LS5 compared to the healthy average (36 species) ^30^. Among these species, *Bifidobacterium longum* climaxed to seven times the HeAve. Finally, there were also isolated blooms of the phyla Fusobacteria (40x and 11x HeAve at LS1 and LS6) and Verrucomicrobia (seven times the HeAve at LS5).

To assess the diversity within each microbial community model, we calculated the alpha diversity based on the AGORA2 taxonomic assignments for both LS1-7 and HeAve. The highest LS alpha diversity was obtained for LS6 (Table 1). Although LS2 was the time point with the highest number of species, it was not the one with the highest alpha diversity, when the taxonomic differences of the different samples were weighted with a hierarchical tree based on the taxonomies ^34^. Indeed, LS2 was mainly composed of Firmicutes and Actinobacteria, which covered more than 70% of the relative abundance in the sample (**Fig. 1**, Table 1, Supplementary Table S1).

**Table I.** Information about the species filtering performed through AGORA2 mapping. The total number of strains identified, and the strains covered by the AGORA2 mapping are reported. The abundance based on AGORA2 mapping, using a cutoff threshold abundance of 0.0001 is reported as well, together with the alpha diversity of the samples. For details on the calculation of the Alpha diversity, please refer to the method section.

| Features | LS1 | LS2 | LS3 | LS4 | LS5 | LS6 | LS7 | HeAve |
| --- | --- | --- | --- | --- | --- | --- | --- | --- |
| **Total number of strains identified** | 1,041 | 1,055 | 1,532 | 1,110 | 1,037 | 1,112 | 1,039 | 939 |
| **Strains covered in the *in silico* microbial community models** | 76 | 125 | 102 | 91 | 87 | 101 | 106 | 198 |
| **Fraction of total abundance covered** | 0.923 | 0.937 | 0.902 | 0.904 | 0.926 | 0.925 | 0.923 | 0.98 |
| **Alpha diversity** | 74.86 | 77.86 | 76.85 | 77.11 | 74.5 | 80.18 | 77.46 | 75.56 |

To assess the changes in diversity between the time points, we calculated the beta diversity using the Bray-Curtis dissimilarity index ^34^. The average beta diversity between samples was 58.00. The two most dissimilar samples were LS1 and LS2 (84.1%), likeliest reflecting the effect of the antibiotic treatment before LS2 collection. The two lowest beta diversities were between LS3 with LS4, which had a beta diversity of 23.44, and LS5 with LS7 of 35.76. The diversity between LS1 and LS6 was 79.50 (Supplementary Table S1).

In the Principal Component Analysis (PCA) performed on microbial composition and abundances (**Fig. 2B**, Supplementary Fig. S1, S2), the first component accounted for 54.7% of the total variability, while both the second and the third components each accounted for approximately 20% of the total variability. The different PCA components were mainly driven by the differential abundance of two archaeal species, i.e., the already mentioned *M. smithii* ATCC 35061, and *Methanosphaera stadtmanae* DSM 3091 (LSMax = 542x HeAve). Both archaea were more abundant in LS1 and LS6 in comparison to the other time points (**Fig. 2A**, Supplementary Fig. S2, S17). In the PCA cluster plot, LS6 was clearly separated from the other six time points (**Fig. 2B**), consistent with the higher alpha diversity of the LS6 microbiome (Table 1) compared to the other time points.

As aforementioned, the LS1 gut microbiome was severely depleted in almost all the most abundant HeAve species (**Fig.1a**, Supplementary Fig. S3-S9). We found 21 microbe species with relative abundance >1% in HeAve, yet they were very rare (HeAve/LS1>10) in LS1 (**Fig. 1B,** Supplementary Fig. S3A), including the phylum Bacteroidetes species *Prevotella copri* (1436x), *Bacteroides stercoris* (152x), *Bacteroides caccae* (43x), *Bacteroides ovatus* (40x), *Bacteroides vulgatus* (28x), *Bacteroides dorei* (20x), *Alistipes putredinis* (15x) and the phylum Firmicutes species *Eubacterium rectale* (43x) and *Ruminococcus bromii* (11x). Only three of the 21 HeAve most abundant species had relative abundances in LS1 that were comparable (1<HeAve/LS1<5) to HeAve: *Faecalibacterium prausnitzii* (4.5x), and *Alistipes finegoldii* (3x), *Dialister invisus* (1.5x). We note that *F. prausnitzii* is a well-known anti-inflammatory bacterium. Its high level at LS1 may be an indication of the microbiome attempting to counter the high level of host inflammation at LS1.

A complementary analysis identified microbial species with the highest relative abundance (>1%) in the gut microbiome of LS1 (Supplementary Fig. S3B). Not only were there fewer microbe species that had a relative abundance >1% than in HeAve, but also the most abundant microbes in LS1 were normally extremely rare in the healthy gut microbiome. With the exception of *Dialister invisus* (which we will return to later), all of the dominant LS1 microbiome species ranged from 100 to almost 1,000 times more abundant than in the healthy gut microbiome. A number of these normally rare species (e.g., *E. coli, M. smithii, M. stadtmanae, and P. micra)* will have major impacts on key metabolite production, as we will discuss later in this paper. This illustrates a classic ecological dynamics result: when formerly dominant species are wiped out, normally rarer species can bloom and become the dominant ones.

Taken together, our metagenomic time series demonstrates that the microbial composition varied substantially between the LS time points, as well as compared to the healthy average. The strong differences between LS and healthy samples in microbial abundance motivated the use of metabolic modelling to understand how these metagenomic differences over time could influence metabolite production as the gut microbiome ecology shifts.

### Microbial and metabolic changes over time

To investigate potential changes in metabolic activity associated with the dysbiotic microbiome composition at each time point, we performed metabolic modelling and FBA ^16^ assuming a Western diet ^35^. For each metabolite, we computed the net metabolite production potential (Methods, Supplementary Table S2). The resulting *in silico* metabolite production profiles represent the potential of all microbial community members to uptake dietary metabolites and secrete metabolic end products. We also predicted microbe-specific contributions to the overall fluxes in each microbial community model (Methods). To allow for the comparison between the LS microbiome and the healthy gut microbiome, we calculated the parameter healthy average of the fluxes (HeAveFluxes) from the healthy controls metagenomic input to our simulation model, creating 34 healthy controls microbial community models (Supplementary Table S2). The resulting HeAveFluxes parameter enabled us to discover that over 20 metabolites had a maximum value over the LS1-7 LS dysbiotic gut microbiome, which ranged from 10 to 750 times higher than the maximum values across the healthy controls (Supplementary Table S2).

Next, we examined in detail the strong time variations of a number of key gut microbially produced metabolites. Specifically, we selected 24 metabolite exchange reactions with LSMax/LSMin flux ratios >10 [or LSMin=0, so the ratio is large (technically divided by 0)] to examine in more depth (Supplementary Table in Supplementary Materials). All but five of these 24 were greatly overproduced by LS, with LSMax from 5x to 750x times the highest value found across the 34 healthy controls. For each of these 24 metabolites, we then visually pattern-matched the metabolite time graph to microbial species relative abundance graphs over time. This approach allowed the identification of several microbe-metabolite relationship time variation patterns over LS1-7 (**Fig. 3,** Supplementary Fig. S15-S21). The microbe-metabolite relationships were characterised by two distinct microbe/metabolite classes: Class I, with a peak value at LS1 and dramatically lower values in the other time points (e.g., the *M. smithii/methane*, **Fig. 3**, Supplementary Fig. S17), and Class II, which were low in LS1 and higher values in the subsequent time points (e.g., *Dorea longicatena* DSM 13814/Pyridoxal, **Fig. 3**). We give archetypal examples of each Class in Figure 3 with subclasses of Classes I and II defined with matching metabolite examples in Supplementary Fig. S15-S21.

Numbers in parentheses next to each metabolite name corresponds to the flux ratio of the minimum and maximum flux calculated for the LS samples. For insights on subclasses of Classes I and II, please refer to Supplementary Fig. S15-S21.

These two dozen microbe species-metabolite pairs were all candidates for a deeper biochemical pathway analysis to determine whether there is a clear causal relationship between the microbe and the production of its paired metabolite. Because of our discovery of the extreme overproduction in the disease state compared to the inter-population variability in healthy individuals, these metabolites are all potential candidates to be biomarkers for tracking the episodic development of the disease. Below, we take a first look at this hypothesis.

#### Methane and *Methanobacteriaceae*

The average methane production in healthy gut microbiomes (HeAveFluxes) computed by AGORA2 was 0.26 mmol/gDW/day (Supplementary Table S2). In contrast, in the LS diseased state, the production of methane was highest in LS1 (40.19 mmol/gDW/day) and decreased in LS2-7 (1 mmol/gDW/day at LS2) closely following *Methanobacteriaceae* abundance in the corresponding microbiome samples (**Fig. 3**, top). This relationship was also reflected in the microbe-metabolite simulations (Supplementary Table S3). At its peak, *M. smithii* was the most abundant species in LS1 (Supplementary Fig. 3B) and methane production in the disease state was 155x the highest value computed (HeMax) for methane across the healthy controls. This result agrees with prior results that methanogenic archaea are the major biological source of methane in humans with a single species, *M. smithii*, accounting for up to 94% of methanogenic activity in most colonised individuals ^36^. In addition, chorismate followed the same time evolution as methane, peaking at 141x HeMax.

#### Oxygen and *E. coli*

The *in silico* average healthy level (HeAveFluxes) of the oxygen production fluxes was 0.003 mmol/gDW/day. In contrast, the extreme value of LS1 was over 2,000 times higher (8.3 mmol/gDW/day) than HeAve and 165x HeMax (we note only two of the 34 healthy controls had any significant oxygen production). This enormous increase in the dysbiotic production of free oxygen was found to follow the time variation of *E. coli*, being highest at L1-L3 (where *E. coli* was ^~^10% of the gut microbiome ecology or 187x HeAve relative abundance), normal at LS5 and LS7, and an additional increase at LS6. It is remarkable how large the change in oxygen production was as the dysbiotic evolution progressed. The ratio of the oxygen production from its high in LS1 (8.3) to its low in LS7 (0.00075 or 0.25x HeAve) was over 10,000 fold (11,116x).

These large fluctuations suggest an obligate syntrophy with one of the oxygen-producing bacteria present in the consortium. Many reconstructions of microbial species included in the simulations (e.g., *Eggerthella lenta* DSM 2243, *Bacteroides vulgatus* ATCC 8482, and *Megasphaera elsdenii* DSM 20460) have a superoxide dismutase (VMH ID: SPODM) converting reactive oxygen species to oxygen and oxygen peroxide (2.0 h[c] + 2.0 o2s[c] -> h2o2[c] + o2[c]), as well as an oxygen exchange (VMH ID: EX_o2) reaction. The simulations, therefore, suggest a novel, unidirectional interaction among these species boosting *E. coli* bloom with superoxide dismutase products. Our hypothesis is novel, yet consistent with the “oxygen hypothesis” that posits that some aspects of IBD symptoms result from an increase of oxygen and reactive oxygen species into the intestinal lumen competitively favouring facultative anaerobic species over strictly anaerobic ones ^37^. Indeed, Enterobacteriaceae bacteria, such as *E. coli*, can absorb oxygen being facultative aerobic species ^38^.

In addition to the likely increased *anaerobic* respiration, which was induced by inflammation in LS (Fig. 1) ^38^, the dysbiotic shifts in the microbiome ecology itself produced, according to our microbial community model, copious amounts of free oxygen, coming from the detoxification of reactive oxygen species, which *E. coli* could then utilise to increase its relative abundance directly via *aerobic* respiration. In addition to the production of free oxygen, our microbial community model predicted that the fluxes of trimethylamine N-oxide (TMAO), for LS1 was 3082x HeAveFluxes (Supplementary Table S2). In previous studies, TMAO has been highlighted as a metabolite, which alters systemic homoeostasis and participates in the first inflammatory states ^39^. Furthermore, TMAO production is known to boost anaerobic respiration, which favours Enterobacteriaceae, i.e., *E.coli*, over Clostridia and Bacteroides species ^40^. Therefore, we conclude that there appear to be two separate mechanisms (inflammation-induced aerobic respiration and dysbiotic microbiome ecology creating free oxygen) that both provide *E. coli* with a selective energy advantage over the otherwise dominant Firmicutes and Bacteroides, which can do neither anaerobic nor aerobic respiration.

#### Thiamine and *E. coli*

Vitamin synthesis by gut microbes is one of their essential ecological services to the health of the host. Our microbial community model predicted that, in the extreme of the disease state (LS1), thiamine (vitamin B1) fluxes (17.1mmol/gDW/day in LS1) was 18,318x higher than the HeAveFluxes (0.00093 mmol/gDW/day) (Supplementary Table S2). The thiamine production flux computed by the LS microbial community models was highly variable, fluctuating across LS1-7 by a factor of 15,000x, while across the population of healthy patients, each sampled at one time, there was a variation in thiamine production of only 23x. Furthermore, the maximum value of thiamine (at LS1) was 7,472x greater than HeMax for thiamine (Supplementary Table S2), meaning that the disease state drove thiamine production almost four orders of magnitude beyond what was seen in the cross-population production.

In addition, we also predicted other B vitamins to be overproduced in LS compared to HeAve (Supplementary Table S2). In particular, our microbial community model predicted LSMax/HeAve for riboflavin (vitamin B2, 8x), pyridoxal (vitamin B6, 98x), and folic acid (vitamin B9, 39x). For niacinamide (vitamin B3) and biotin (vitamin B7), the healthy controls were all zero, but there was substantial production of each in LS1-7. This result highlights the value of measuring the dysbiotic time variation within a single patient instead of only reporting population averages.

#### Other metabolites that vary with *E. coli*

Several other metabolite exchange fluxes (**Fig. 3**, Supplementary Fig. S18), which closely followed the time variation in *E. coli* relative abundance, were also found to have an LSMax value far above cross-population HeMax value. Specifically, the ratio of LSMax/HeMax for some of these included the polyamine metabolism-related metabolites ortho-hydroxyphenylacetic acid (164x), 5’methylthioadenosine (142x), spermidine (12x), and histamine (13x) consistent with previously reported results ^41^. In particular, dysbiosis can predispose overgrowth of *E. coli*, which in turn leads to increased production of histamine, thus contributing to the symptomatology of histamine intolerance ^42^. The LS1-7 variation predicted for histamine was 170x, with a maximum flux of 46.95 mmol/gDW/day in LS1, while the HeAve was 0.32 mmol/gDW/day (Supplementary Table S2).

#### TMAO and Fusobacterium species

TMAO has been recently hypothesised to be a possible link mediating between red meat intake and vascular inflammation, leading to poor cardiometabolic health ^43^. Separately, Fusobacteria have been discussed as being involved in the onset of colon cancer ^44^. Intriguingly, at the height of LS inflammation, as measured by calprotectin and serum CRP, LS1 had nearly a 1,000x overabundance of the dominant phylum Fusobacteria species *Fusobacterium sp. 12_1B*, compared to HeAve. This coincided with our microbial community model predicting a similar level of overproduction of TMAO (LS1/HeAve=3082x) and LS1/HeMax=91x.

#### Serotonin and *B. longum*

The microbial production of the neurotransmitter serotonin, which had a variation across LS1-7 of 25x and whose maximum at LS5 was 3x the HeMax, mimicked the fluctuations of *Bifidobacterium longum* abundances (Supplementary Fig. S19). It is known that *B. longum* supernatants upregulate the serotonin transporter expression in intestinal epithelial cells ^45^. Deregulation of gut-produced serotonin has also been associated with diarrhoea or constipation symptoms ^46^. Furthermore, according to Minderhaud and colleagues ^47^, the severity of intestinal inflammation can depend on the availability of gut serotonin. Another metabolite, which followed the variation of *B. longum* and is also involved in the gut-brain axis, was tryptamine, which varied by 103x across LS1-7 and whose peak at LS1 was 25x HeMax.

#### *Dorea longicatena* and *Ruminococcus obeum*

were the 1st and 2nd most abundant Firmicutes species in the Lachnospiraceae family in LS, respectively. Their time evolution aligned with a number of metabolites with large ratios of LSMax/HeMax: formaldehyde (3477x), 5-methyltetrahydrofolic acid (311x), tetrahydrofolic acid (260x), folic acid (12x), pyridoxal (7x), and riboflavin (3x).

#### Normally rare Firmicutes species

An unusual aspect of LS dysbiosis was that at LS1 several normally quite rare Firmicutes species *(Peptostreptococcus stomatis*, LS’s most abundant species in Firmicutes family Peptostreptococcaceae; *Solobacterium moorei*, LS’s most abundant species in Firmicutes family Erysipelotrichaceae; and *Parvimonas micra*, LS’s most abundant species in Firmicutes family Clostridiales Family XI. Incertae Sedis) were from 250 to 1,000x more abundant than HeAve. Their graph over LS1-7 closely matches metabolites with high ratios of LSMax/HeMax: acetoin (38x), trimethylamine (13x), and 1,2-diacyl-sn-glycerol (8x) (Supplementary Fig. S15).

Taken together, we observed that dozens of metabolites were greatly overproduced by LS in the disease state compared to our healthy controls and we discovered numerous microbe-metabolite pairs showing similar changes over time. Overall, our analyses at the different time points as well as longitudinally illustrate that the dysbiotic microbial composition changes were associated with significant changes in metabolic function.

### Metabolic and subsystem signature of each phase

After analysing metabolites strongly diverging between LS and healthy average patients, we focused on reactions subsystem and metabolites characterising the different phases of the disease development. The constraint-based modelling approach revealed that the reaction subsystems strongly changed during the disease progress, which was associated with changes in metabolic production potential by the microbial communities (Supplementary Table S2). It has been reported that the prevalence or absence of reaction subsystems in microbial community models can reflect healthy or dysbiotic microbial communities ^48^.

First, we investigated which metabolite production potentials followed the observed proximity of LS3 with LS4, and LS5 with LS7 in the PCA plot (**Fig. 2A**). The net production of some metabolites increased or decreased constantly from LS3 to LS5 but were predicted to be very high (isobutyrate) or very low in LS6 (**Fig. 4**). The net flux production of L-isoleucine, ethanol, and L-lactate was low in LS1 and LS6 (HeAveFluxes 130.48 mmol/gDW/day), while it increased in the other time points (HeAveFluxes 234.93 mmol/gDW/day). In contrast, the production of isobutyrate followed an opposite trend and had a higher simulated accumulation in LS1 and LS6 (HeAveFluxes 106.87 mmol/gDW/day) compared to the other samples (HeAveFluxes 30.14 mmol/gDW/day). With the exception of isobutyrate, each of the metabolites in Figure 4 has a ratio of their maximum values in LS1-7 (LSMax) that were greater than the maximum value (HeMax) in any of the 34 healthy individuals: butyrate (2.7x), ethanol (2.6x), L-isoleucine (7.3x), and L-lactate (3x). These results suggest that monitoring the fluctuations of key microbial species and key metabolites together with the biological processes of bioconversion could help to identify transitions of inflammation (**Fig. 4**).

**Fig. 4.**
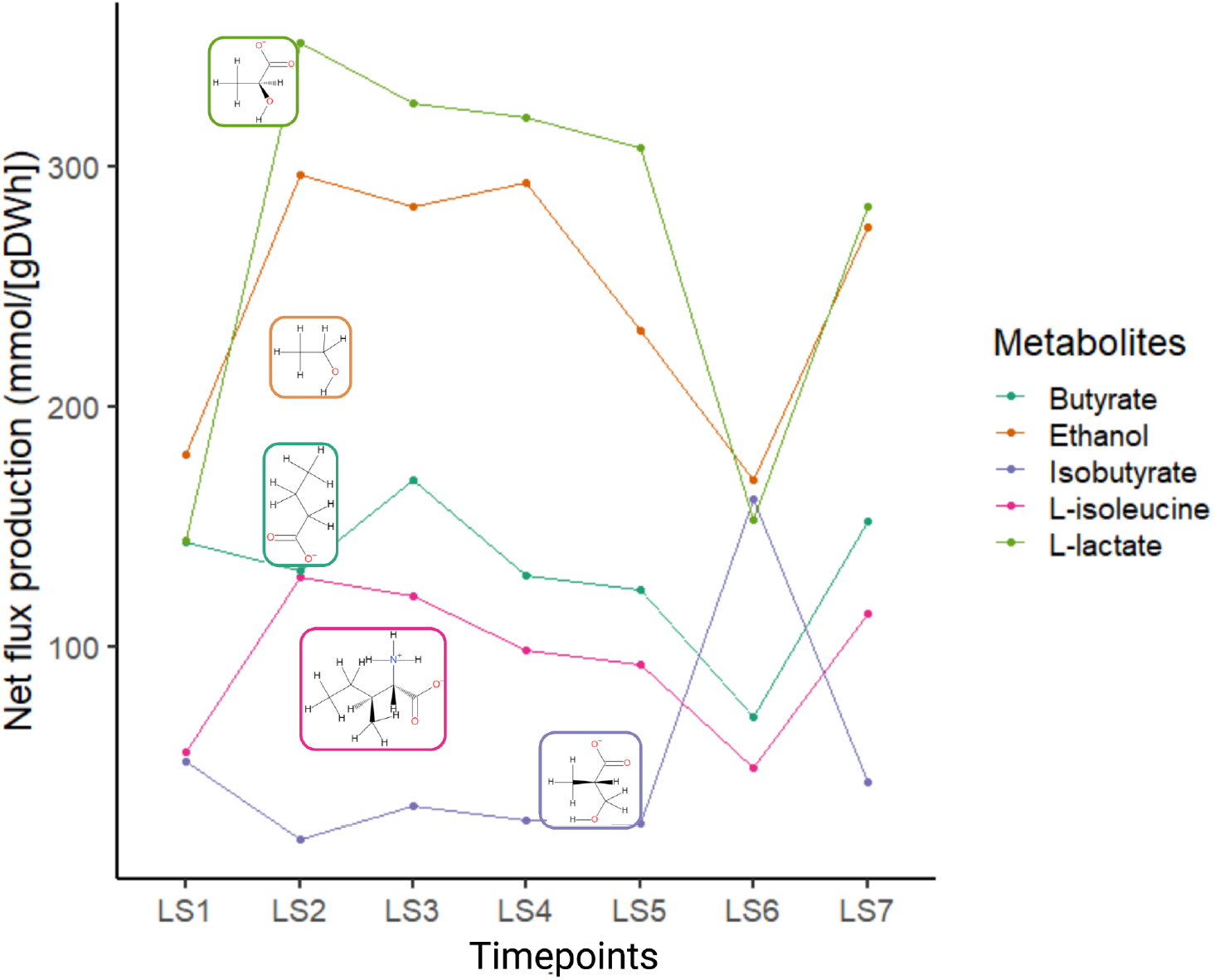
Metabolites net flux variation. Line plot of the net flux production (mmol/gDWh) of metabolites having a marked change over the different time points. For each metabolite, the respective chemical structure is reported.

To identify metabolite signatures at each time point, Euclidean clustering ^49^ was performed considering all the metabolites predicted with net flux production higher than ten mmol/gDW/day. The results revealed the existence of three main clusters (**Fig. 5A**). The threshold of ten was selected arbitrarily for graphical purposes. The first cluster, named low fluxes (LF), grouped together all the metabolites with a very low net production; the second cluster had the metabolites with intermediate net production (IF); the third included the metabolites with high net production (HF). The three clusters were heterogeneous in metabolic subsystem composition. Some metabolites, whose flux rates were variable among the different phases of the disease, will be discussed more in detail and the roles of the microbial species mainly involved in their production. The prevalence of subsystems including reactions related to that metabolite will be discussed as well.

**Figure 5:**
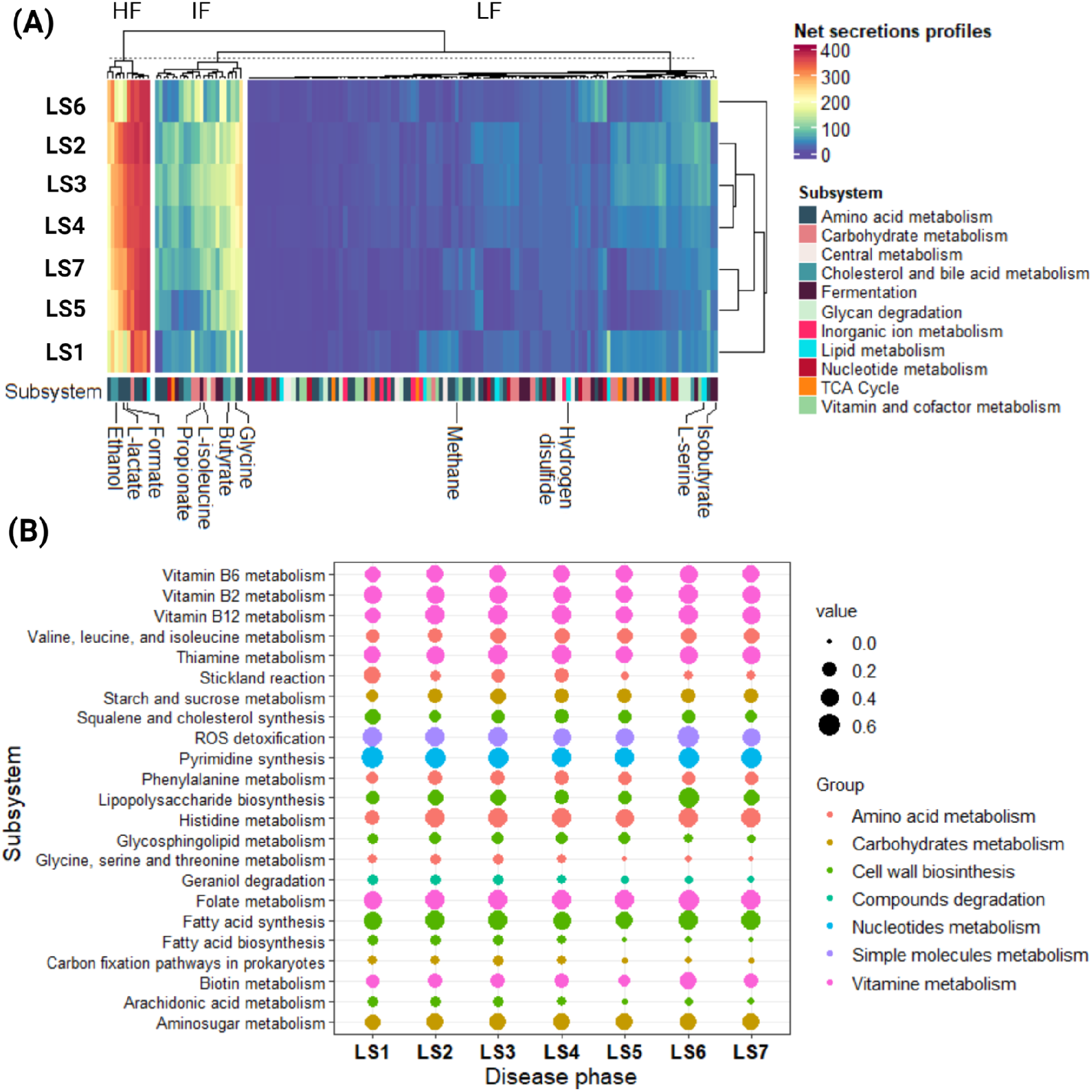
Overview of metabolites produced and reactions subsystems across the different time points. **(A)**Heatmap of the net flux production of all metabolites with a summed net flux higher than ten mmol/gDW/day. Key metabolites commented on in the text have been highlighted in the heatmap. LF - Low flux; IF - Intermediate flux; HF - High flux (see text for more information). **(B)**Geom plot of reaction subsystem prevalence across the different time points. The colours of the circles refer to the “manually-attributed” group of each subsystem. The diameter of the circles is proportional to the abundance of the reactions in the modelled microbial communities.

As expected, “Methane metabolism” was strongly increased in LS1 and LS6 compared to the other phases (Supplementary Table S2) and was due to the higher abundance of methanogenic Archaea in LS1 and LS6. Accordingly, the predicted production of methane enhanced in LS1 and LS6 (log fold change (LogFC) 1.42 and 0.84, respectively, Supplementary Table S2). In contrast, some subsystems were phase-specific (Supplementary Table S2). This was the case for the “Stickland reaction” (**Fig. 5B**), which couples oxidation and reduction of amino acids to organic acids ^50^ and characterised LS1. In a study exploring the subproducts of common degradation pathways, 80% (8/10) of Stickland reaction products have been frequently detected in IBD patient stool ^51^. Since all microbial community models for the seven time points received the same *in silico* diet, the observed differences were a direct result of the difference in microbial composition. The increase in hydrogen production could be an additional cause of the bloating event experienced as one of the IBD symptoms. Levitt and Olsson have already linked hydrogen production to the adverse bloating event ^52^. LS6 was characterised by an increased abundance of *E. coli* strains, e.g., *E. coli* 042, *E. coli* B354, and *E. coli* FVEC1302, whose genomes encode enzymes belonging to lipopolysaccharide (LPS) biosynthesis subsystems. LPSs are produced and secreted by gram-negative bacteria (e.g., *Salmonella typhimurium* ^53^ and *E. coli* ^54^), and can provoke an immune response. LPSs are generally soluble as monomers but they can aggregate into fibrous and highly insoluble lipoproteins and lead to inflammation ^55^. It has been reported that the concentration of LPS is increased in the acute phases of the disease compared to relapsing ones ^56^.

Butyrate is a key energy source for the host’s colonic epithelial cells ^57^. Our microbial community models predicted that the butyrate secretion rate in LS7 (flux of 152.164 mmol/gDW/day) was more than twice as high as the LS6 butyrate secretion rate (70.58 mmol/gDW/day). This jump was confirmed by the laboratory-measured butyrate concentration in the LS6 and LS7 faecal samples (Supplementary Table S1, Fig. S14), which also more than doubled from 0.7 to 1.7 mg/mL. This large increase in butyrate production is likely driven by the 8-fold increase, from 3.99% (LS6) to 31.5% (LS7) (**Fig. 3**, Supplementary Table S1), of the relative abundance of *Faecalibacterium prausnitzii*, one of the major butyrate-producing microbes in the human gut.

Furthermore, the production of L-serine was increased in LS6 (**Fig. 3**) compared to the other time points (Supplementary Table S2). L-serine has been shown to interact with the gut microbiome, in particular, it is known to elicit the secretion of antimicrobial molecules, such as bacteriocins ^58^. This amino acid is mainly produced by species of the *Dialister* genus and its importance will be discussed in the following paragraph. *E. coli* pathogenic strains have been proposed to use L-serine anabolism to enhance their fitness in the inflamed gut ^59^. In contrast, this pathway has a minor role in the pathogenic bacterial growth of healthy guts ^60^, suggesting that the signals or transduction pathways necessary for L-serine catabolism activation could be responsible for pathogen-specific adaptation to the inflammatory microenvironment. Intestinal inflammation can result in the generation of a microenvironment that is conducive to the growth of Enterobacteriaceae, allowing the outcompete of obligate anaerobes ^61^. Therefore, enterobacterial blooms, such as those seen in LS (Supplementary Fig. S10), and more generally in CD, are a hallmark of inflammation-associated dysbiosis ^62^. Accordingly, *E.coli* can catabolize L-serine converting it to pyruvate, a crucial substrate for gluconeogenesis and tricarboxylic acid cycle pathways ^63^. L-serine also plays a role as a signalling molecule targeting the expression of stress response genes ^58^. Furthermore, it can be used as a precursor in the synthesis of gene products involved in stress adaptation ^59^. In this context, it is known that L-serine catabolism is increased in *E. coli* under heat shock conditions and L-serine is used for the generation of heat shock proteins ^59^. L-serine uptake during inflammatory conditions is probably a conserved mechanism utilised by pathogenic bacteria for their competitive fitness ^64^.

Taken together, our results revealed that numerous metabolite production fluxes were altered during the seven time points. However, further validation in other patients or hypothesis testing in model organisms will be required.

### Insight into *Dialister spp*. metabolism and net of interactions

Next, we aimed at elucidating which microbes were driving the metabolic changes at each phase, thereby, shedding light onto the potential mechanisms of the disease onset. Therefore, we calculated the microbe-metabolite contribution using the cooperative tradeoff algorithm ^65^. Briefly, this algorithm assumes that the growth rate of an individual microbe in the community is maximised, while a sub-optimal growth rate of the remaining microbial community is maintained. The microbe-metabolite contribution identified that in LS6, the production of L-serine was mediated mainly by two members of the *Dialister* genus, i.e., *Dialister succinatiphilus* YIT 11850 (Max (LS1-7) = 19x HeAve) and *Dialister invisus* DSM 15470 (Max (LS1-7) = 0.9x HeAve).

*D. invisus* DSM 15470 has been shown to be involved in the establishment of dysbiosis typical for the IBD gut microbiome ^66^. Hence, metabolites produced by species of this genus were inspected in detail. *D. invisus* DSM 15470 was identified in all time points except LS5, but its metabolic activity, measured in terms of the number of exchanged compounds, was very different in these time points. In LS1, *D. invisus* produced as many as 12 metabolites and consumed 52, while in the other time points, it consumed an average of ten metabolites. Notably, *D. invisus* and *D. succinatiphilus* YIT 11850 produced L-serine and formate during LS1, and glycine in the other phases (**Fig. 6**, Supplementary Table S3). Dietary glycine is known to prevent chemical-induced colitis by inhibiting the induction of inflammatory cytokines and chemokines ^67^.

**Figure 6:**
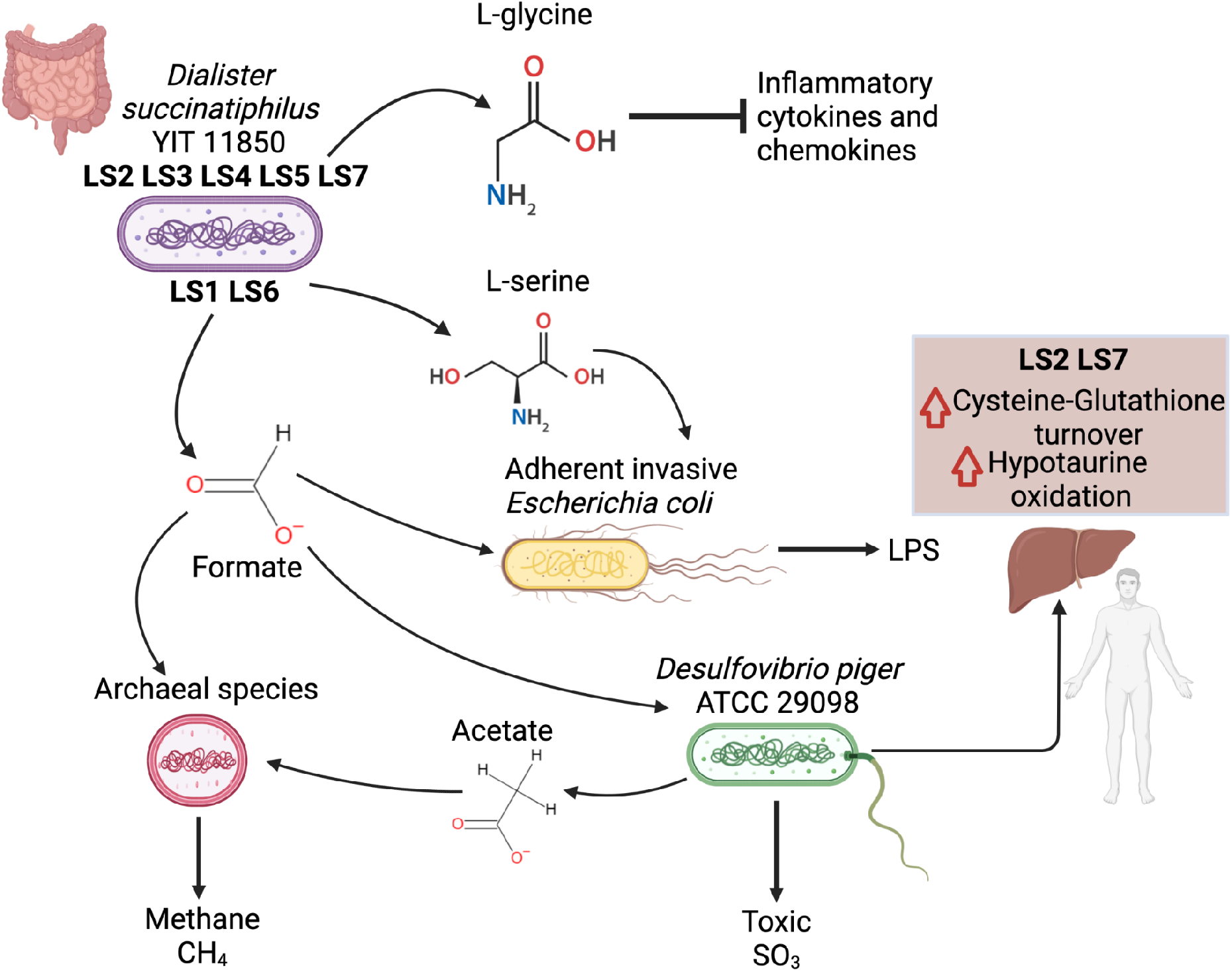
Map of species interactions. The map shows the most relevant microbial species and their interconnection in a dependency network. The figure was created with BioRender.

Both L-serine and formic acid have been proposed to mediate proinflammatory mechanisms ^60^. In LS6, L-serine uptake was mainly mediated by members of the Enterobacteriaceae family (e.g., *E. coli* 042, B354, FVEC1302, and H299). Additionally, formate production in LS1 was assigned a key role in the microbial interaction (**Fig. 6**). Indeed, formate in LS1 fuelled the proliferation of *E. coli* F11, *M. smithii* ATCC 35061, and *D. piger* ATCC 29098 (**Fig. 6,**Supplementary Table S3). *E. coli* F11 is an adherent invasive and pathogenic strain, which takes advantage of the leaking gut to replace strictly anaerobic bacteria ^68^. *M. smithii* ATCC 35061 is a hydrogenotrophic archaeon that can use either CO2 and H2 or formate alone for methane production ^36^. The increase in methane production, and, therefore, constipation and bloating events, is known to be partially caused by the increase of this archaeal species (**Fig. 6,** Supplementary Table S3) ^69^. The prevalence of *D. piger* is higher in patients hospitalised for IBD in comparison to healthy individuals or patients hospitalised for other pathologies ^70^.

### The multifaceted role of *Desulfovibrio piger* ATTC2

The microbial composition varies between individuals ^71^, which may not necessarily translate into functional or metabolic differences ^72^. However, certain metabolic functions may require the presence of specific microbial species ^73^. Hence, we investigated whether there were any function-specific microbes in the microbial community models at the different time points, whose presence was required for the production of specific metabolites. The analysis revealed that the Proteobacteria *D. piger* ATCC2 was the only microbial species involved in the production of sulphite (SO_3_^2-^) in the microbial community models (Supplementary Table S3). We found that D. piger at its peak in LS was nearly four times more abundant than in the maximum relative abundance found across the healthy controls (LSMax/HeMax=3.7). Patients affected by IBD, such as ulcerative colitis, are strongly discouraged to consume foods with high SO_3_^2-^ levels as being harmful and favouring tightening of inflammation ^71^. Furthermore, sodium sulfite, a common food additive, inhibits the activity of commensal and anti-inflammatory bacteria, such as *F. prausnitzii* ^73^. A large part of the SO_3_^2-^ present in the gut comes from dietary intake ^74^, however, some microbial species are known to produce SO_3_^2-^. *D. piger* ATTC2 was not able to synthesise SO_3_^2-^ in single-species simulations. However, the pairwise simulations revealed that this species interacted with the Archaea *M. stadtmanae* DSM3091. Only when in synergy with the archaeal partner, *D. piger* ATTC2 could produce SO_3_. This metabolic dependency reflected a cooperative behaviour culminating in the production of the host-toxic SO_3_^2-^. To support this hypothesis, it is worth noting that the flux production of SO_3_^2-^ in the LS time points, strongly reflects the abundance fluctuations of both *M. stadtmanae* and *M. smithii*, the two archaea in the community under investigation (Supplementary Fig. S17).

The pairwise simulations revealed that *D. piger* ATCC 29098 absorbed ethanol, converted it into acetate, and, finally, secreted it. The acetate was then taken up by the acetoclastic Archaea *M. stadtmanae* DSM 3091. *D. piger* ATCC 2909 can metabolise ethanol using two alternative anaerobic pathways: in one case, the ethanol is oxidised to acetate via acetaldehyde as an intermediate. In the second case, other intermediates between acetaldehyde and acetate are generated, namely acetyl-CoA and acetyl-P ^75^. In the simulations, the conversion of ethanol to acetate had a yield of approximately 1 (0.93), as expected from experimental data ^76^. The nearly 1:1 ethanol-to-acetate ratio reflected the release of an excess of reducing equivalents, such as methane, by syntrophic partners ^75^. *D. piger* ATCC 29098 ability to synthesise and export SO3 may be recovered through the interaction with other archaeal partners as well. Accordingly, due to the commensalism, when *M. stadtmanae* DSM 3091 and *D. piger* ATCC 29098 were co-occurrent, the net flux production of methane was higher (Supplementary Table S2). The abundance of *D. piger* strains in IBD patients has already been reported ^70^.

Taken together, our results indicate that through the production of two metabolites, i.e., L-serine and formate, species of the *Dialister* genus cooperate with many pathogenic strains, such as adherent invasive *E. coli* strains, archaeal species, and *D. piger* ATCC2. The interactions could trigger inflammatory responses and enhance methane production. Furthermore, *D. piger* ATCC2 plays an important role in enhancing the production of host-toxic SO3^2-^ in microbial communities.

### Whole-body modelling suggests a role of *D. piger* in the transsulfuration pathway

We then integrated the microbial community metabolic models of each time point with a male organ-resolved whole-body model of human metabolism ^27^ to track the metabolic consequences of gut microbiome dysbiosis on different body sites, organs, and tissues on the host metabolism. In this simulation, we inspected the microbial metabolic influence on a range of different organs and tissues of the host as the only variable was the gut microbiome ecology composition, which changed over time. We found that the dysbiosis resulted in greater flux changes in some organs or cell types than in others. In fact, red blood cells, platelets, and the retina showed the most pronounced changes in predicted fluxes (**Fig. 7A**, Supplementary Table S4). Interestingly, Episcleritis, a disease involving the eyes, is one of the most common extraintestinal IBD manifestations ^77^. The predicted flux through the metabolism of the prostaglandin E2 was altered in the pancreas in LS2 and LS7. Prostaglandin release is one of the first triggering factors of the inflammatory cascade typical of CD ^78^. Finally, many drugs, such as mesalazine, are used to inhibit the release of prostaglandins and leukotrienes in different body sites ^79^ underlying the key role of these molecules in the establishment of CD.

**Figure 7:**
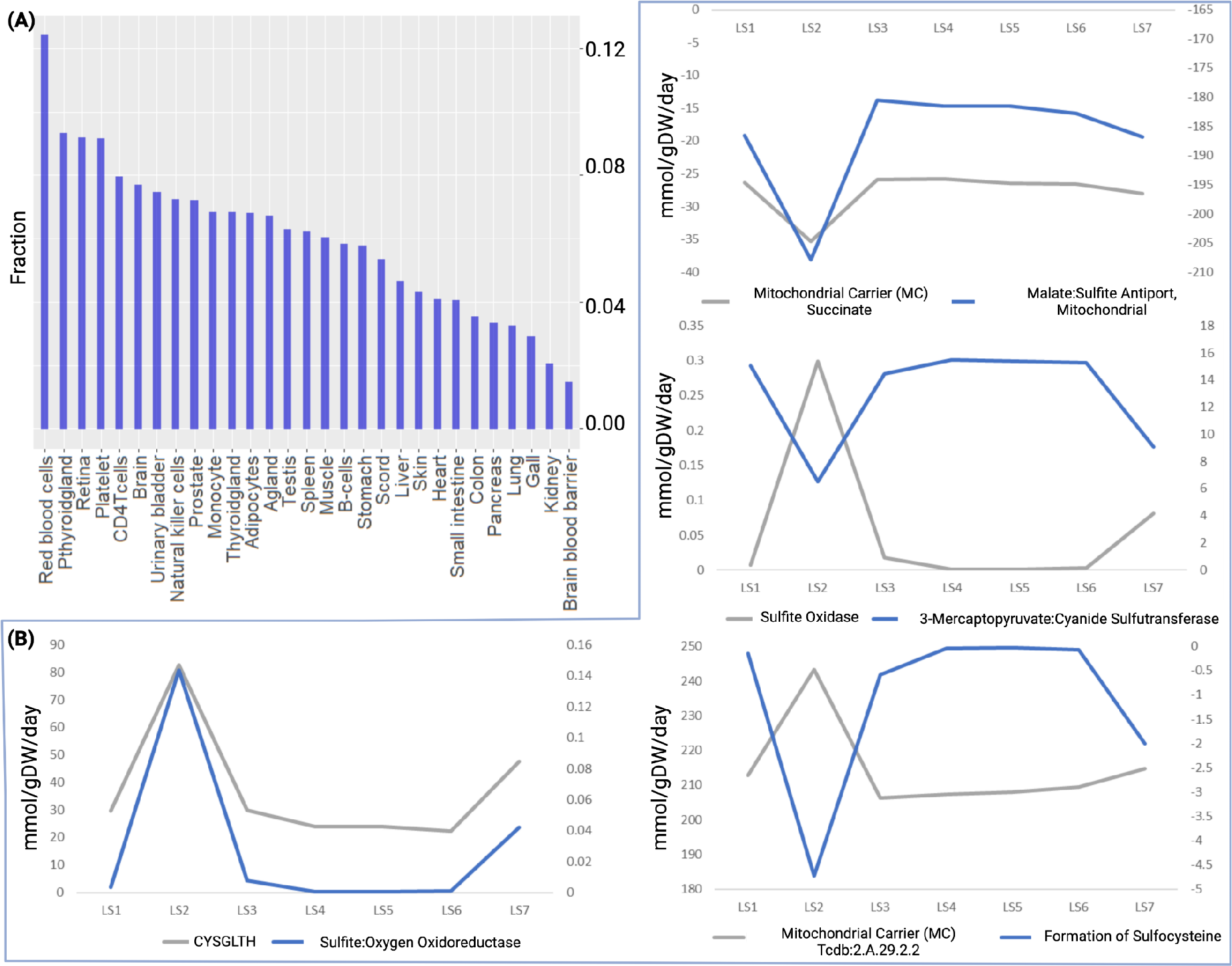
Overview of the alteration of the microbiome-associated whole-body models. **(A)** Fraction of reactions changing flux direction due to the dysbiosis divided by body sites. **(B)**Key reactions in the sulphur metabolism in the liver are altered by dysbiosis (Supplementary Table S4).

Notably, the sulphite metabolism in the liver was dependent on the presence of *D. piger* ATCC 2909 (**Fig. 7,** Supplementary Table S4) in the gut. In the microbiome-associated whole-body model, sulphite from the small and large intestine could be either transported directly to the liver through the portal vein or as cysteine-S-sulphate (VMH ID: slfcys). In the liver, cysteine-S-sulphate can then be metabolised to cysteine (VMH ID: cys_L) and SO_3_^2-^ ^80^. In our simulations, SO_3_^2-^ was oxidised to SO_4_^2-^ through sulphite oxidase activity (VMH ID: SULFOX). This reaction produced hydrogen peroxide in the model and thus could contribute to oxidative stress, which we do not model as such. At the same time, sulphur metabolism was linked to the metabolism of bile acids in the simulations in LS2 and LS7. Indeed, cysteine-S-sulphate is a precursor of hypotaurine and taurine. The flux through the reactions involved in this pathway (involving the reactions HYPTROX, r0539, and r0381) changed with the time points, being increased in LS2 and LS7. Around 85% of bile acid reactions were affected by the presence of *D. piger* ATCC 2909 (**Fig. 7B**, Supplementary Table S4). The increase of these compounds led to an increase in faecal H_2_S, which has been reported to be increased in IBD patients compared to healthy controls ^81^, in LS2 and LS7. Cysteine was converted to L-cystine (Supplementary Fig. S20) by a state transition reaction named Glutathione:Cystine Oxidoreductase (VMH ID: CYSGLTH). CYSGLTH enabled the oxidation of glutathione; thus, acting as a scavenger molecule. Based on our predictions, the presence of *D. piger* changed the fluxes in the transsulfuration pathway leading to higher cysteine to glutathione turnover.

In conclusion, the analysis of the microbiome-associated whole-body model has the potential to shed light on the alteration in the metabolism of different body sites caused by the dynamic dysbiotic microbiome. The alteration of the sulphur metabolism in the liver and its link with the presence or absence of *D. piger* ATCC 2909 in the large intestine reflects the intercommunication among the different body sites. This network and its influence on the disease onset has been so far largely overlooked. However, this connection could reveal understudied pathobiology mechanisms.

## Conclusions

The time course analysis performed on a patient affected by IBD enabled the analysis of how net metabolite production fluxes were induced by the change of the gut microbial community composition during a time course covering more than one year. To date, this is the first analysis inspecting the disease evolution in a patient affected by Crohn’s disease with metabolic modelling. The study revealed that substantial metabolic changes are associated with the disease evolution as a direct consequence of the patient’s changing gut microbiome composition, notably involving archaeal species. A number of biologically important metabolites were found to be highly overproduced over time in the patient, compared to healthy controls. This list includes oxygen, methane, thiamine, formaldehyde, TMAO, folic acid, serotonin, histamine, and tryptamine, which may yield new biomarkers of disease progression. The analyses with microbiome-associated whole-body models revealed that the presence of *D.piger* could alter the metabolism of sulphur in the liver. Since the microbial composition is variable among individuals, to obtain a wide and general representation of the microbiome the time course inspection of a higher number of patients will be needed. However, the functional redundancy present in the gut microbiota allows to some extent to generalise our results to other patients. Taken together, we demonstrated that microbial community metabolic modelling is a very valuable *in silico* approach to track correspondence between metagenomic data and metabolite production and can yield testable novel hypotheses to be addressed with additional validation studies. These results underline the importance of tracking an individual’s gut microbiome composition and metabolic production along a time course, paving the way to new analyses for self-quantified medicine.

## Methods

### Ethics statement

The stool samples of the patient were collected by consent under two protocols: HRPP 141853 (American Gut Project) and HRPP 150275 (Evaluating the Human Microbiome). The protocols include written informed consent concerning dissemination and scientific publication of the results. Both protocols were approved by the Human Research Protection Program (HRPP) of the University of California, San Diego.

### Longitudinal sample collection

The samples have been collected from naturally passed faeces and immediately stored without a buffer at −80°C. Seven samples were selected. A personal symptom log entry has been generated at the time that each faecal sample was passed. Additionally, the weight and body mass index (BMI) of the patient have been determined on the day associated with each sample. Other metadata were obtained from previous publications ^22^.

### Metagenomics data generation

Metagenomic sequencing of the seven stool samples (LS1-LS7) was performed at the J Craig Venter Institute using Illumina HiSeq2000 platform. On average, 160 million paired-end reads at 2×100 base pairs were generated per sample. The raw reads for the healthy controls were downloaded from National Center for Biotechnology Information (NCBI) Sequence Read Archives under BioProject ID PRJNA43021. The processing of the raw metagenomic sequence data for LS and for the healthy controls and the computation of species relative abundance were described in an earlier publication by Wu et al ^28^. Briefly, after low-quality reads, reads from humans and duplicated reads were removed, and the filtered reads were then aligned to our curated microbial genomic sequences. The reads were assigned to their top matched genomes and the depth of genome coverage of each species was calculated and then normalised to relative abundance so that the total relative abundance was 1.0.

### Butyrate and biomarkers measurement

The patient LS used the company Doctor’s Data Comprehensive Stool Analysis kit (www.doctorsdata.com/Comprehensive-Stool-Analysis-CSA21) to generate the values of lysozyme, lactoferrin, and secretory IgA (all ELISA) reported in Figure 1. In addition, total butyrate (Supplementary Material Figure S14) was measured from the LS stool sample by Doctor’s Data using gas chromatography. The values in Figure 1 of faecal calprotectin, generated by ARUP Laboratories using Immunoassay, and of serum CRP were from tests with UC San Diego Health.

### Definition of the average European diet

The *in silico* diet represented the nutrient intake of an average European individual, hence, representing a typical “Western” diet. Its description, along with the corresponding flux values, was obtained from the nutrition resource in the Virtual Metabolic Human database ^35^. The diet was supplemented with metabolites that have been previously ^24^ determined as necessary for the biomass production of at least one AGORA reconstruction. The dedicated function (adaptVMHDietToAGORA.m) of the Microbiome Modelling Toolbox ^82^ was used to constrain each microbial community model. The lower bounds on all other dietary exchange reactions were set to zero to prevent the uptake of other metabolites.

### Simulations

Simulations were carried out using the COBRA Toolbox ^82^ and the Microbiome Modelling Toolbox ^83^, which is part of the COBRA Toolbox, in MATLAB version 2018b (Mathworks, Inc.) as a programming environment. Metagenomics reads were mapped onto the AGORA2 collection to create the microbial community models for the simulation. For this purpose, the function translateMetagenome2AGORA from the COBRA Toolbox was used. Microbial species with relative abundance higher than 10^-5^ were considered in the population analysis (i.e., for the alpha and beta analysis), while for the AGORA2 collection mapping and all microbial community models, a threshold of 10^-4^ was used. The precise total relative abundance covered for each time point is reported in Table 1. Abundances were normalised for the microbial community modelling. For the simulations and the net secretion and uptake fluxes predictions, the function initMgPipe, contained in the Microbiome Modelling Toolbox ^84^, was used. More specifically, the function initMgPipe contains the function microbiotaModelSimulator, which calculates the net maximal production capability for each metabolite. This parameter indicates the maximal production of each metabolite and is computed by summing the maximal secretion flux with the maximal uptake flux for each metabolite. Furthermore, the function initMgPipe contains the function adaptVMHDietToAGORA, which was used to apply the diet constraints to the microbial community model. Microbe-metabolite contributions were performed following Basile et al ^13^. In brief, the MICOM software ^65^ was used through the cooperative trade-off algorithm integrating the abundances as input. Subsystems had been assigned following the procedure proposed by Heirendt et al. ^83^, and implementing the function calculateSubsystemAbundance using as input the reaction abundances.

The integration of the whole-body model was performed using the Harvey reconstruction ^27^. To create the personalised gut model, the function combineHarveyMicrotiota was used and the simulations were performed with the minNorm algorithm through the COBRA Toolbox (optimizeWBmodel).

For all simulations, the optimisation solver used was CPLEX (IBM iLOG, Inc).

### Statistical analysis

A cohort of 34 metagenomic samples from 34 healthy individuals from the Human Microbiome Project ^32^ was used to create a “healthy average” (HE Ave) value for each microbe species. Then, we computed the ratio of the relative abundance of the seven time points to the average health and reported the ratio of the maximum value at any of the seven time points, i.e., Max (LS1-7), to the healthy average. Alpha diversity and beta diversity analysis were calculated with the “vegan” package ^34^ and using R software v.4.0.3. The taxonomic differences of the different samples were weighted with a hierarchical tree based on the taxonomies of AGORA2 ^25^ with the function taxa2dist. The alpha diversity was calculated with taxondive ^85^. The score considered for the alpha diversity was Δ*. For the beta diversity, the function vegdist was applied. The values of beta diversity were converted to Newick format and used to generate a tree representing the differences between samples with the function NJ of the ape package. The PCA was performed with the function princomp with the parameters “cor=TRUE, scores=TRUE”. The 3D plot of the PCA was realised with the function plot3d of the package “rgl”. The Log2 Fold Change was adopted as a parameter to characterise metabolite production across samples.

## Data availability statement

The authors are in the process of submitting the metagenomic sequences of the specific sequencing done at JCVI for the LS1-7 samples.

Two later publications resequenced some of the LS1-7 samples, at a lower depth than reported herein, as part of research on a longer time series of LS faecal samples. The first publication ^22^ resequenced LS 1-7 (12/28/2011 to 4/29/2013) as part of a longer time series of 27 LS samples (dates from 12/28/2011 to 12/07/2014 are listed in their Supplementary Table S1) analysing the metagenomics of E. coli strain dynamics. The metagenomics sequence of these 27 samples can be found in EBI under study PRJEB24161. The second publication ^30^ sequenced eight LS time series samples (dates from 12/28/2011 to 5/22/2016), including resequencing LS1-3, and added metaproteomic analysis for these eight time points. Metagenomic data are available through EBI under the study PRJEB28712 (ERP110957).

## Supplementary Material

**Supplementary Table S1:**

- Metadata of the different time points. The table accounts for the collection date, the age of the patient, the clinical signs, as well as the bmi and the blood concentrations of some markers (i.e. CRP, calprotectin, lactoferrin, lysozyme, iga)
- Details of the raw reads used for this manuscript including number of raw reads (pair), QC filtered reads (pair), reads without human sequences or duplicates (pair), reads mapped to reference genomes (pair)
- Relative abundances of microbial species mapped on AGORA2 in the different LS samples, the maximum value found in LS, the Healthy average and their ratio are reported as well
- Relative abundances of microbial strains in the different LS samples retrieved with metagenomics, the maximum value found in LS, the Healthy average and their ratio are reported as well
- Relative abundances of microbial species in the different LS samples retrieved with metagenomics, the maximum value found in LS, the Healthy average and their ratio are reported as well
- Relative abundances of microbial strains mapped on AGORA2 in the healthy samples
- Relative abundances of microbial strains in the healthy samples retrieved with metagenomics
- Relative abundances of microbial species in the healthy samples retrieved with metagenomics
- Details on taxonomy of the species retrieved from metagenomics accounting for all the taxonomic levels available and the taxid
- Phyla relative abundances for HE Ave and LS1-7
- Calculation of the alpha diversity of the different microbiomes
- Calculation of the beta diversity between the different time points

**Supplementary Table S2:**

- Abundance of the reactions presence/absence for each time point considered, HeAve is reported
- Abundance of the subsystems presence/absence for each time point considered, HeAve is reported
- Abundance of the subsystems presence/absence for all the healthy patients considered
- Net secretion of simulated metabolites for each LS time point considered, average net secreted fluxes from Healthy patients are reported as well
- Net secretion of simulated metabolites for each healthy patient considered
- Log fold change of simulated metabolites for each time point considered

**Supplementary Table S3:**

- Microbial metabolite contribution simulated for all the different time points

**Supplementary Table S4:**

- Fluxes simulated for all the time points considered integrating the whole-body model and the microbiome information
- Information on all the reactions considered including description, formula, and crosslinks to other databases

**Supplementary Material:**

- Supplementary Figure S1: Scree plot of the PCA in Figure 2 of the manuscript
- Supplementary Figure S2: Rotating PCA accounting for the three space dimensions better describing the variance observed in the LS samples
- Medical history of LS
- Supplementary Figure S3A: The 21 microbe species with a relative abundance >1% in the HeAve microbiome (blue bars). For each species the red bar shows the relative abundance in sample LS1. Note that almost all normally abundant species in healthy individuals are severely reduced in LS1. For instance, the two most abundant species in healthy individuals, *Bacteroides vulgatus* and *B. ovatus* have values LS1/HeAve of 0.03x and 0.017x respectively
- Supplementary Figure S3B: The 11 species in LS1 that have relative abundance >1% compared to their relative abundance in HeAve. Note that LS1 has blooms of HeAve rare species, such as *M. smithii* (165x HeAve), with overabundance ratios for other species as great as ~1,000x
- Supplementary Figure S4: The 14 species in LS2 that have relative abundance >1% compared to their relative abundance in HeAve. Note that LS2-4, *E. coli* is ~180x HeAve, while *Collinsella aerofaciens* peaks at 55x HeAve in LS2
- Supplementary Figure S5: The 19 species in LS3 that have relative abundance >1% compared to their relative abundance in HeAve. Note that *Dorea longicatena* and *[Ruminococcus] obeum* are 10-20x HeAve in LS3 and 4
- Supplementary Figure S6: The 17 species in LS4 that have relative abundance >1% compared to those species relative abundance in HeAve. Note that *Streptococcus thermophilus* [Firmicutes Class Bacilli] peaks at ~150x in LS4
- Supplementary Figure S7: The 12 species in LS5 that have relative abundance >1% compared to those species relative abundance in HeAve. Note that *Bifidobacterium animalis* [Phylum Actinobacteria] peaks at over 1500x HeAve in LS5
- Supplementary Figure S8: The 22 species in LS6 that have relative abundance >1% compared to those species relative abundance in HeAve. Note that the 2^nd^ most abundant Archaea *(Methanosphaera stadtmanae)* peaks at ~500x HeAve in LS1 and 6
- Supplementary Figure S9: The 10 species in LS7 that have relative abundance >1% compared to those species relative abundance in HeAve. Note that LS7 *F. prausnitzii*, an anti-inflammatory bacteria has a relative abundance of ~ 1/3 of the microbiome
- Supplementary Figure S10: Abundance fluctuations of the main Enterobacteriaceae bacteria present in the LS samples
- Supplementary Figure S11: Abundance fluctuations of *E. coli* in LS samples and healthy average patients
- Supplementary Figure S12: Abundance fluctuations of microbes of the Fusobacteriaceae family in LS samples and healthy average patients
- Supplementary Figure S13: Abundance fluctuations of microbes of the Methanobacteriaceae family in LS samples and healthy average patients, insight on *Methanosphaera sadtmanae*
- Supplementary Figure S14: Total butyrate measured in the time points covered by this analysis
- Supplementary Table in Supplementary Material: an overview of the 24 most divergent metabolites between LS and Healthy patients. For each of the metabolites, the HeMax, HeMin, HeAve, LSMax, LSMin and informative ratios are reported
- Supplementary Figure S15: Correlations between microbial abundances and specific fluxes, Class IA: High on LS1, Low on LS 2-7
- Supplementary Figure S16: Correlations between microbial abundances and specific fluxes, Class IB: high on LS1, other peak at LS6
- Supplementary Figure S17: Correlations between microbial abundances and specific fluxes, Class IC: High on LS1/2, LS5, LS7
- Supplementary Figure S18: Correlations between microbial abundances and specific fluxes, Class ID: High on LS1, 2, and 3, with Another Peak at LS6, Normal on LS 5 and 7
- Supplementary Figure S19: Correlations between microbial abundances and specific fluxes, Class IE
- Supplementary Figure S20: Correlations between microbial abundances and specific fluxes, Class IIA
- Supplementary Figure S21: Correlations between microbial abundances and specific fluxes, Class IIB: Low on LS1, 5, and 7, High on LS2-4, and LS6

## Acknowledgments

This work was financially supported by the “Budget Integrato della Ricerca Dipartimentale” (BIRD198423) PRID 2019 of the Department of Biology of the University of Padua, entitled “SyMMoBio: inspection of Syntrophies with Metabolic Modelling to optimise Biogas Production” to LT. Furthermore, this study was funded by grants from the European Research Council (ERC) under the European Union’s Horizon 2020 research and innovation programme (grant agreement No 757922), by the National Institute on Aging grants (1RF1AG058942 and 1U19AG063744), and from the Science Foundation Ireland under Grant number 12/RC/2273-P2 to IT. The Ph.D. fellowship of AB was supported by “Progetto di Eccellenza DiBio” of the University of Padua. AB was the recipient of the EMBO short-term fellowship 8720. Larry Smarr thanks the UC San Diego Calit2 Qualcomm Institute and the Center for Microbiome Innovation members for useful discussions and a private donor for financial support for this paper. We thank staff at the J Craig Venter Institute for performing the metagenomic sequencing of stool samples and the San Diego Supercomputer for providing the CPU hours for processing the metagenomic sequencing data. A final acknowledgement to the Italian Consortium for Biotechnologies (CIB) for the support.

## Author contributions

Arianna Basile: Funding acquisition, Conceptualisation, Investigation, Formal Analysis, Visualisation, Writing - Original Draft. Almut Heinken: Software, Methodology, Formal Analysis, Writing - view & Editing. Johannes Hertel: Supervision, Writing - Review & Editing. Larry Smarr: Funding acquisition, Formal Analysis, Writing - Review & Editing. Weizhong Li: Review & Editing. Laura Treu: Supervision, Writing - Review & Editing. Giorgio Valle: Writing - Review & Editing. Stefano Campanaro: Supervision, Conceptualisation, Writing - Review & Editing. Ines Thiele: Conceptualisation, Supervision, Funding acquisition, Software, Writing - Review & Editing.

