## Supplemental Material for "Longitudinal flux balance analyses of a patient with Crohn’s disease highlight microbiome metabolic alterations"

**Medical history of Larry**

In 2011 at the University of California Health System, Dr. William J. Sandborn diagnosed LS, at age 63, with colonic Crohn's disease (CCD). Via colonoscopy and abdominal magnetic resonance imaging (MRI) analysis, the inflamed region of the colon was determined to be confined to 6" to 8" of the sigmoid colon. Specifically, in 2012 the colonoscopy revealed that this region was affected by extensive diverticulosis and inflammatory focal ulceration, inflammatory pseudopolyps, and patchy friability not associated with the diverticular orifices. There was no surgery performed on the patient during the time period covered by this work. Lastly, the main symptoms experienced by LS are rectal bleeding, abdominal cramps, bloating, and malaise.

| Names | HeMax | HeMin | HeAve | LSMax | LSMin | Ratio LSMax/HeAve | Ratio LSMax/MaxHe | Ratio LSMax/LSMin |
|---|---|---|---|---|---|---|---|---|
| Thiamine | 0.0 | 0.0 | 0.0 | 17.1 | 0.0 | 18317.9 | 7471.6 | 15209.0 |
| Formaldehyde | 0.1 | 0.0 | 0.0 | 42.1 | 0.1 | 3476.9 | 405.7 | 372.7 |
| Trimethylamine N-oxide | 0.4 | 0.0 | 0.0 | 40.8 | 0.0 | 3082.4 | 90.7 | #DIV/0! |
| Ortho-Hydroxyphenylacetic | 0.1 | 0.0 | 0.0 | 8.3 | 0.0 | 2604.8 | 164.3 | 11088.4 |
| Oxygen | 0.1 | 0.0 | 0.0 | 8.3 | 0.0 | 2557.4 | 164.7 | 11115.6 |
| Glucose 6-phosphate | 0.2 | 0.0 | 0.0 | 14.1 | 0.0 | 2052.1 | 60.4 | #DIV/0! |
| 5'-Methylthioadenosine | 0.2 | 0.0 | 0.0 | 23.0 | 0.2 | 1466.3 | 142.4 | 105.1 |
| (R)-Acetoin | 3.2 | 0.0 | 0.3 | 122.6 | 4.0 | 487.9 | 38.0 | 31.0 |
| 5-Methyltetrahydrofolic acid | 1.4 | 0.0 | 0.1 | 31.8 | 0.8 | 310.8 | 22.4 | 41.6 |
| Tetrahydrofolic acid | 1.7 | 0.0 | 0.1 | 31.4 | 0.8 | 259.6 | 18.6 | 40.1 |
| Tryptamine | 1.6 | 0.0 | 0.2 | 41.4 | 0.4 | 189.9 | 25.2 | 103.1 |
| Spermidine | 2.4 | 0.0 | 0.2 | 30.1 | 0.9 | 158.4 | 12.4 | 32.3 |
| Methane | 3.0 | 0.0 | 0.3 | 40.2 | 1.0 | 155.2 | 13.6 | 40.2 |
| Histamine | 3.5 | 0.0 | 0.3 | 47.0 | 0.3 | 145.4 | 13.3 | 170.5 |
| Chorismate | 0.0 | 0.0 | 0.0 | 0.0 | 0.0 | 140.7 | 12.8 | 37.9 |
| Pyridoxal | 0.1 | 0.0 | 0.0 | 0.7 | 0.0 | 98.0 | 7.1 | 715.6 |
| Trimethylamine | 3.3 | 0.0 | 0.7 | 41.3 | 3.5 | 61.6 | 12.5 | 11.8 |
| Folic acid | 2.9 | 0.0 | 0.9 | 33.9 | 2.3 | 39.3 | 11.7 | 14.9 |
| 1,2-Diacyl-sn-glycerol | 0.0 | 0.0 | 0.0 | 0.0 | 0.0 | 35.6 | 7.8 | 17.2 |
| Serotonin | 3.5 | 0.0 | 0.3 | 10.0 | 0.4 | 32.3 | 2.8 | 24.9 |
| Riboflavin | 12.1 | 0.0 | 4.4 | 35.4 | 2.8 | 8.0 | 2.9 | 12.5 |
| Ubiquinone-8 | 0.0 | 0.0 | 0.0 | 0.0 | 0.0 | 5.4 | 1.0 | 14.0 |
| Xanthosine | 62.0 | 1.2 | 15.9 | 38.2 | 1.5 | 2.4 | 0.6 | 25.2 |
| Phosphate | 2.3 | 0.3 | 0.8 | 1.0 | 0.0 | 1.2 | 0.4 | 50.8 |

**Supplementary Table** in Supplementary Material: an overview of 24 of the most divergent metabolites between LS and Healthy patients. For each of the metabolites, the HeMax, HeMin, HeAve, LSMax, LSMin and informative ratios are reported

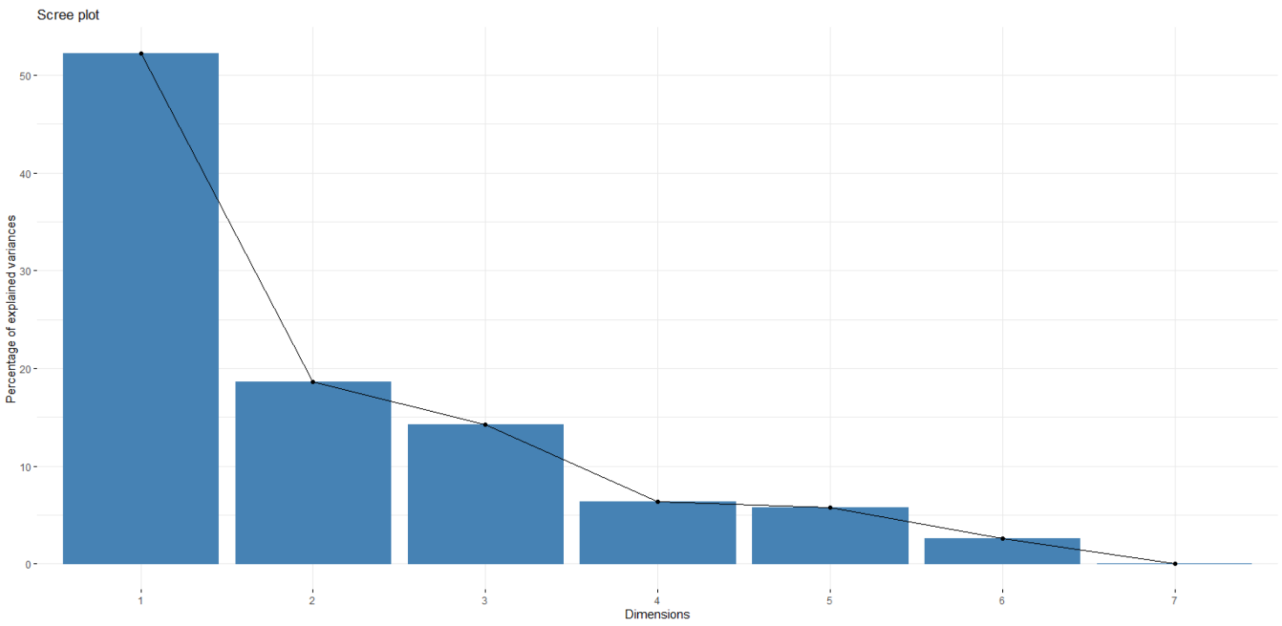


**Supplementary Figure S1:** Screen plot of the PCA in Figure 2 of the manuscript.

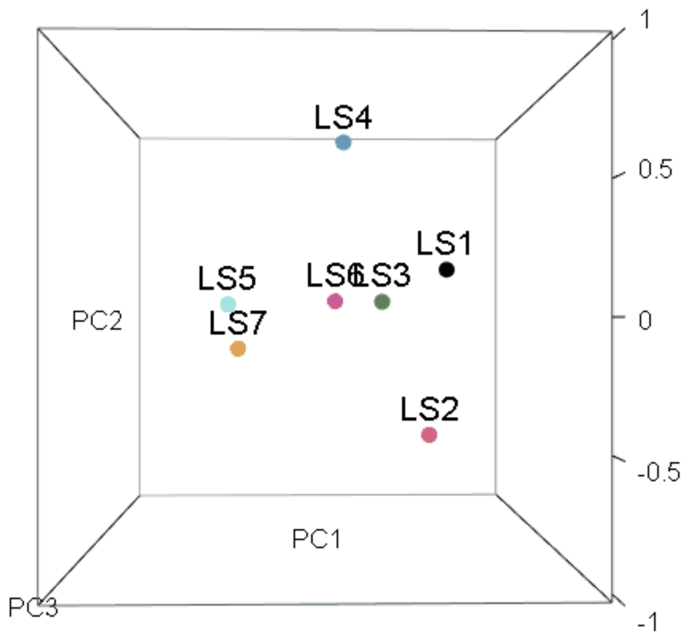


**Supplementay Figure S2:** Rotating PCA accounting for the three space dimensions better describing the variance observed in the LS samples.

**Supplementary Figures 3-9** compare the relative abundance of the cross-population HeAve microbiome with the LS1-7 individual dynamic microbiome. In the graphs blue bars represent HeAve and the red bars represent the LS time samples.

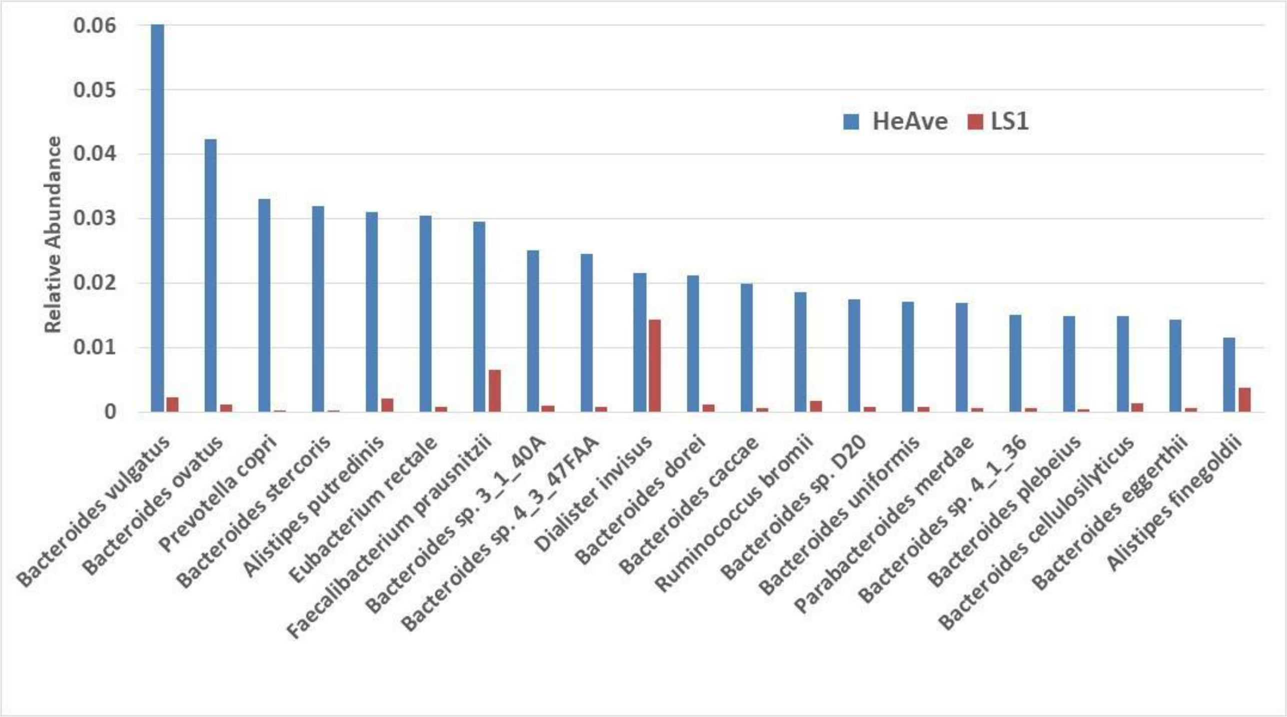


**Supplementary Figure S3A:** The 21 microbe species with a relative abundance >1% in the HeAve microbiome (blue bars). For each species the red bar shows the relative abundance in sample LS1. Note that almost all normally abundant species in healthy individuals are severely reduced in LS1. For instance, the two most abundant species in healthy individuals, Bacteroides vulgatus and B. ovatus have values LS1/HeAve of 0.03x and 0.017x respectively.

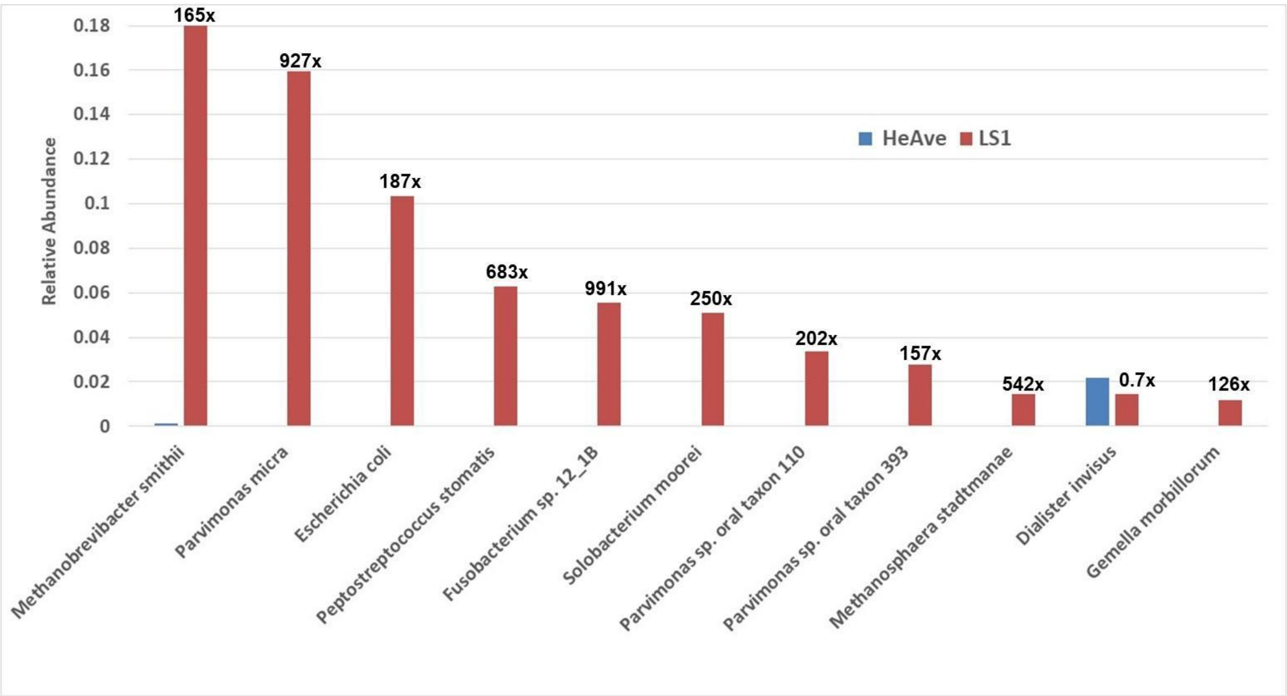


**Supplementary Figure S3B:** The 11 species in LS1 that have relative abundance >1% compared to their relative abundance in HeAve. Note that LS1 has blooms of HeAve rare species, such as M. smithii (165x HeAve), with overabundance ratios for other species from 100 to 1000x.

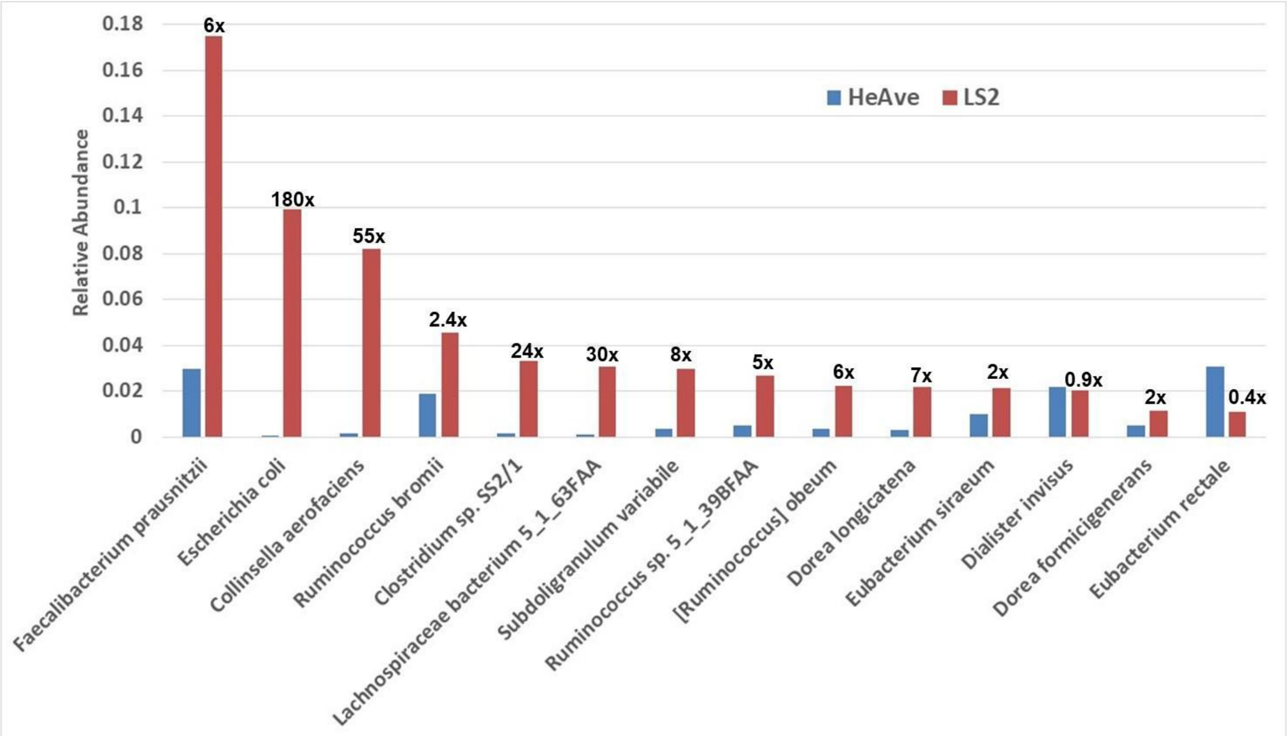


**Supplementary Figure S4:** The 14 species in LS2 that have relative abundance >1% compared to their relative abundance in HeAve. Note that for LS2-4, E. coli is ~180x HeAve, while *Collinsella aerofaciens* peaks at 55x HeAve in LS2.

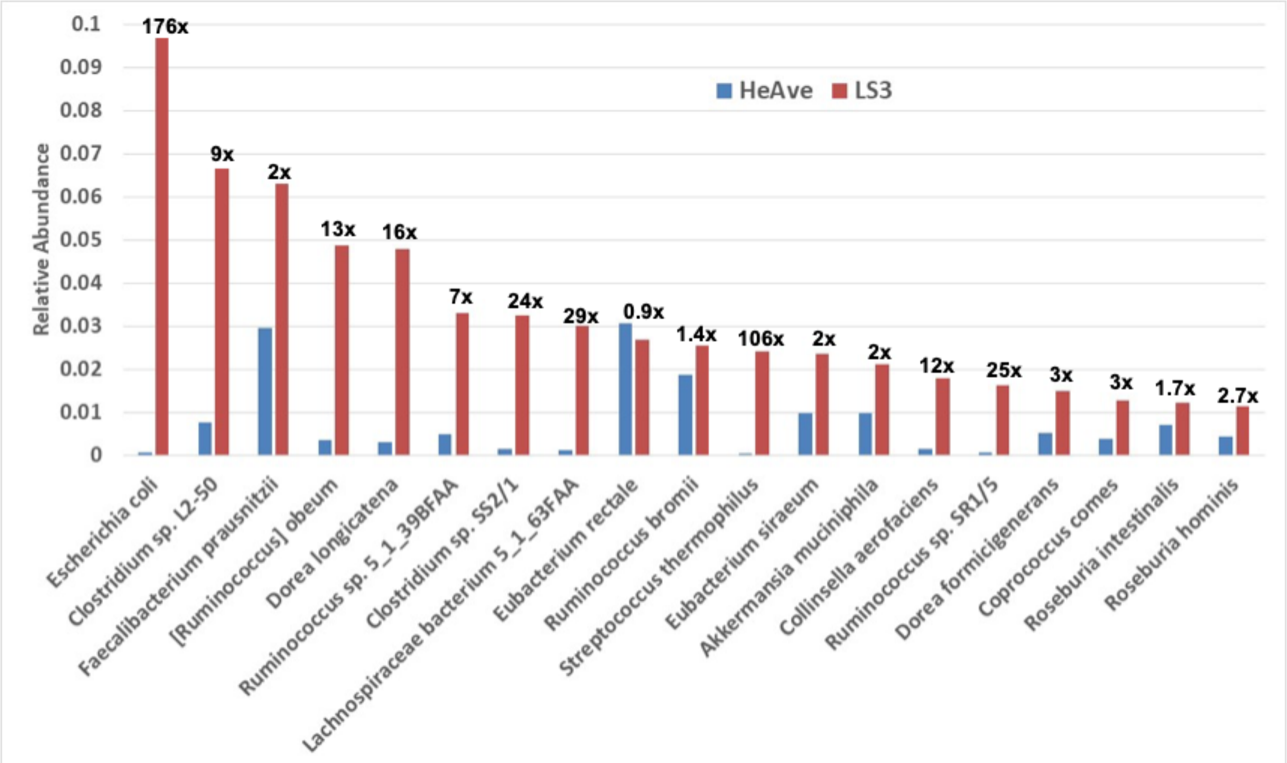


**Supplementary Figure S5:** The 19 species in LS3 that have relative abundance >1% compared to their relative abundance in HeAve. Note that *Dorea longicatena* and [*Ruminococcus*] *obeum* are 10-20x HeAve in LS3&4.

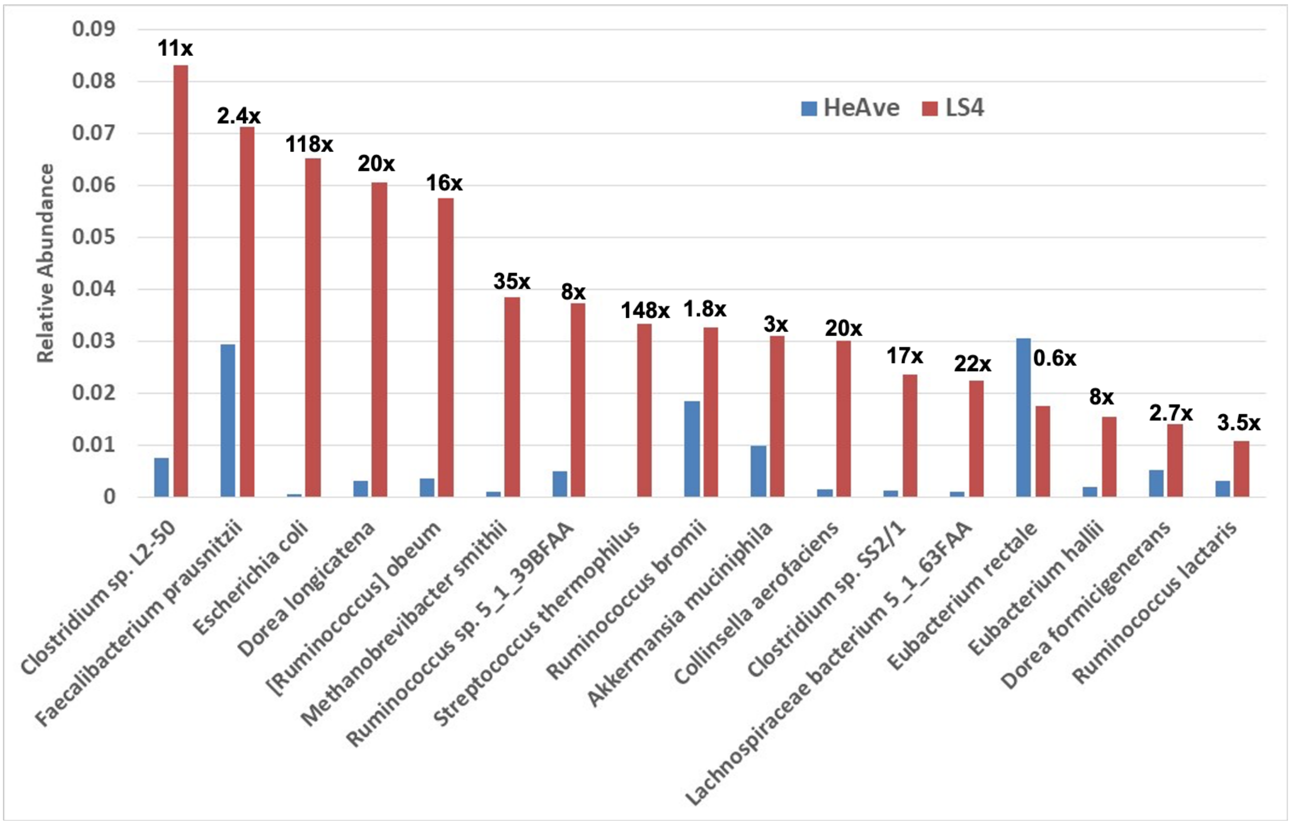


**Supplementary Figure S6:** The 17 species in LS4 that have relative abundance >1% compared to those species relative abundance in HeAve. Note that *Streptococcus thermophilus* [Firmicutes Class Bacilli] peaks at ~150x in LS4.

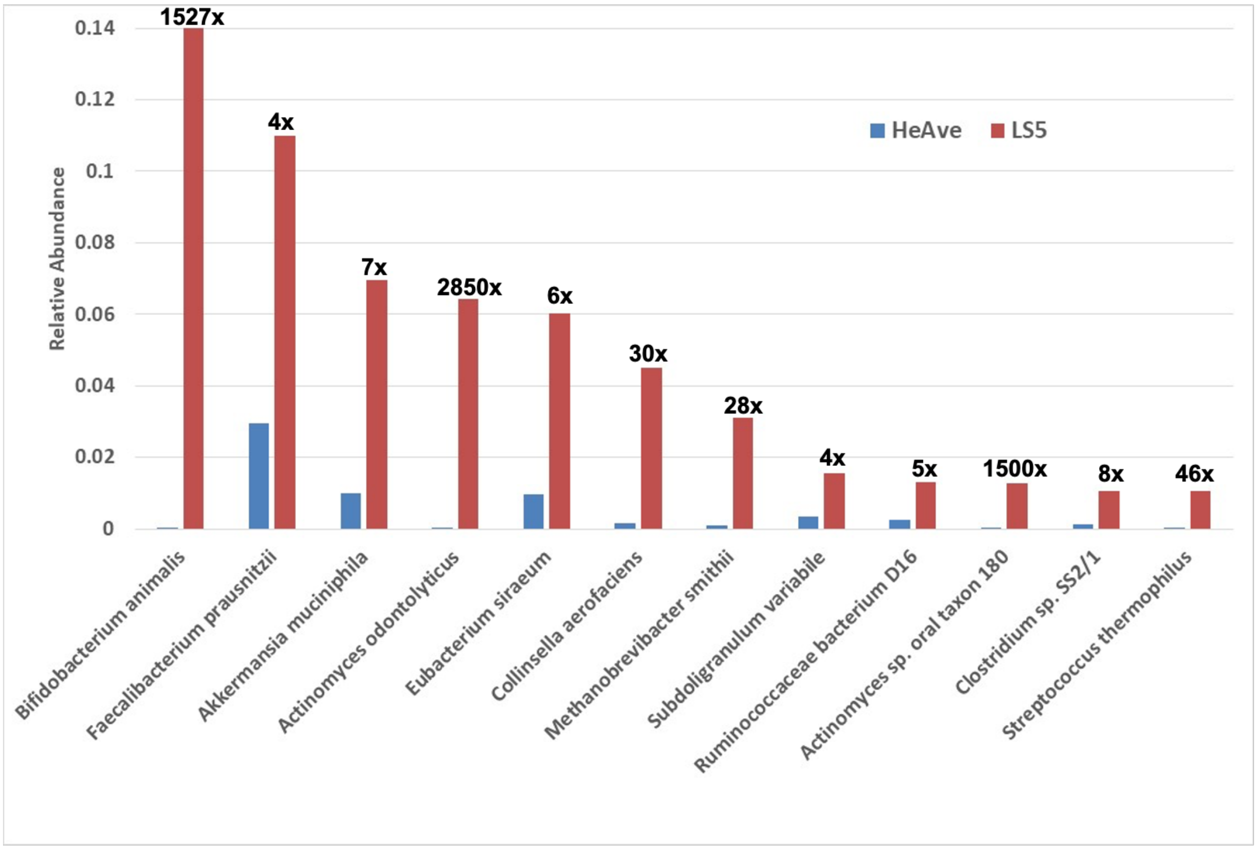


**Supplementary Figure S7:** The 12 species in LS5 that have relative abundance >1% compared to those species relative abundance in HeAve. Note that *Bifidobacterium animalis* [Phylum Actinobacteria] peaks at over 1500x HeAve in LS5.

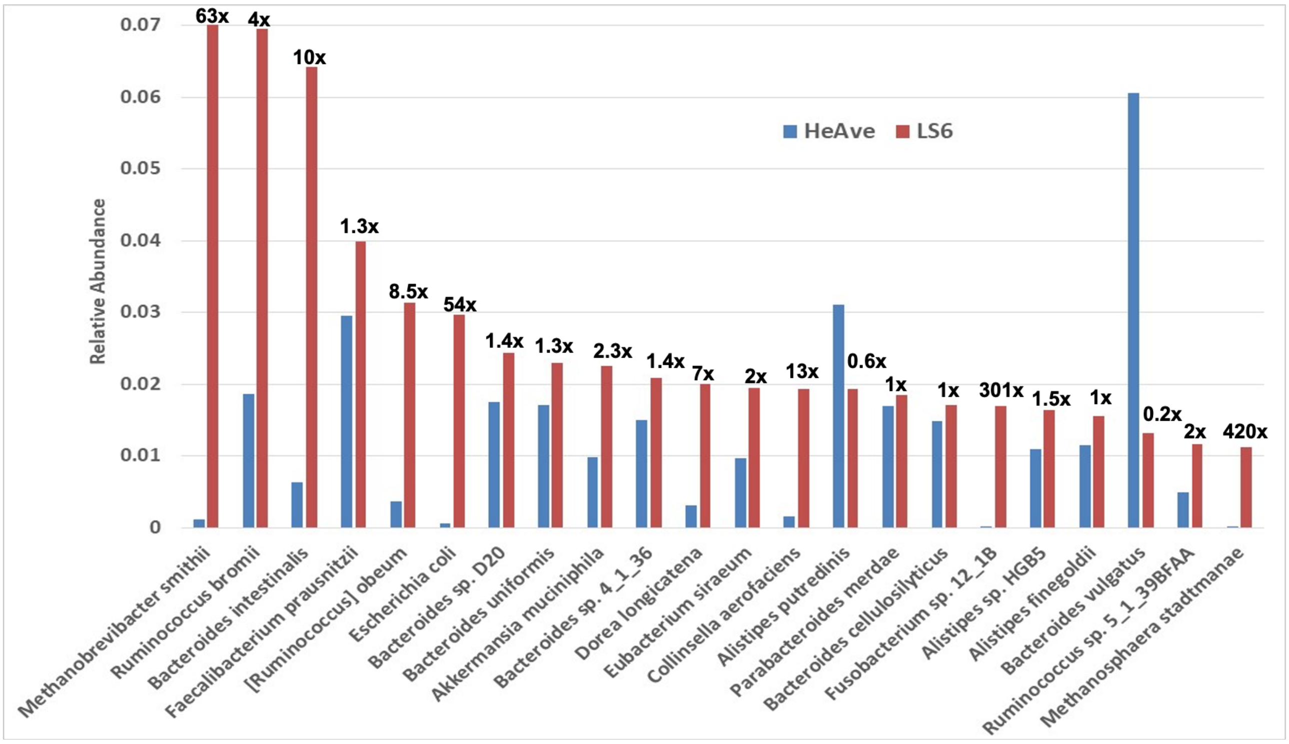

**Supplementary Figure S8:** The 22 species in LS6 that have relative abundance >1% compared to those species relative abundance in HeAve. Note that the 2nd most abundant Archaea (*Methanosphaera stadtmanae*) peaks at ~500x HeAve in LS1&6.

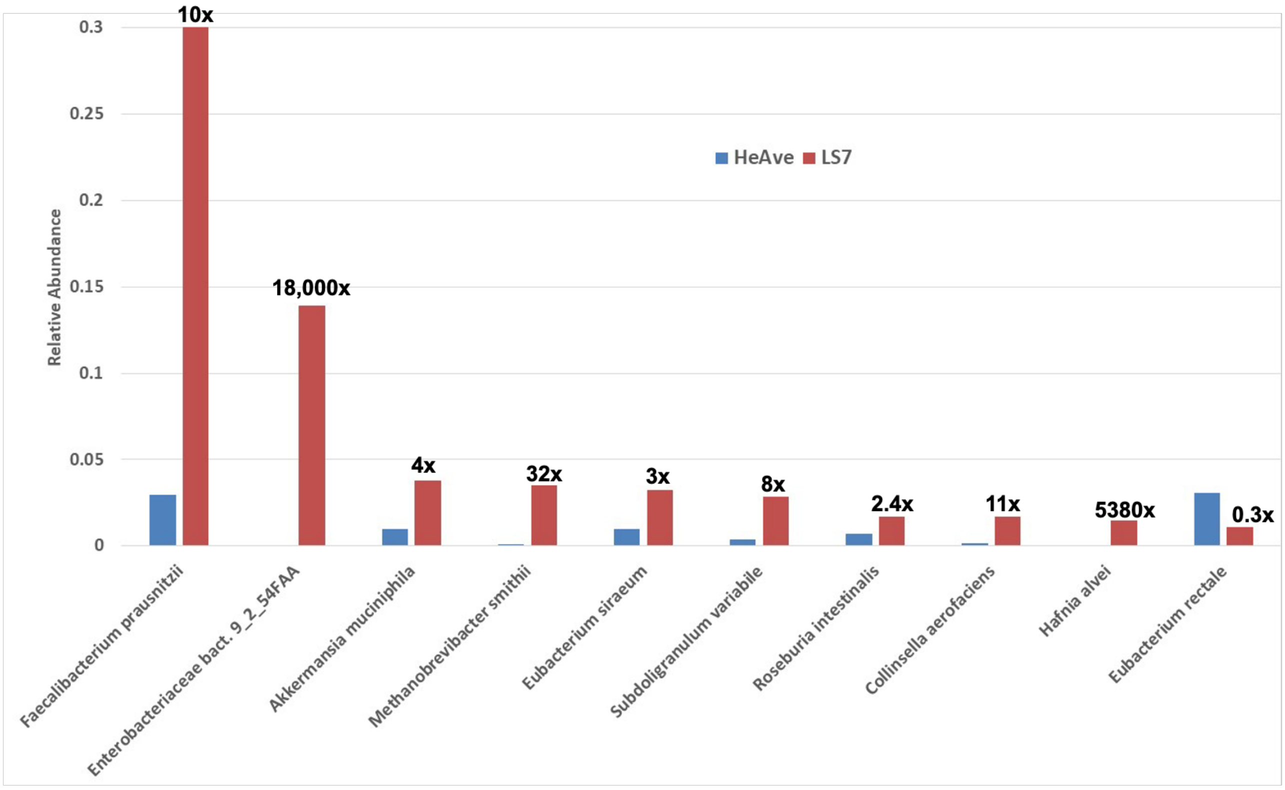


**Supplementary Figure S9:** The 10 species in LS7 that have relative abundance >1% compared to those species relative abundance in HeAve. Note that LS7 *F. prausnitzii*, an anti-inflammatory bacteria, has a relative abundance of ~ 1/3 of the microbiome.

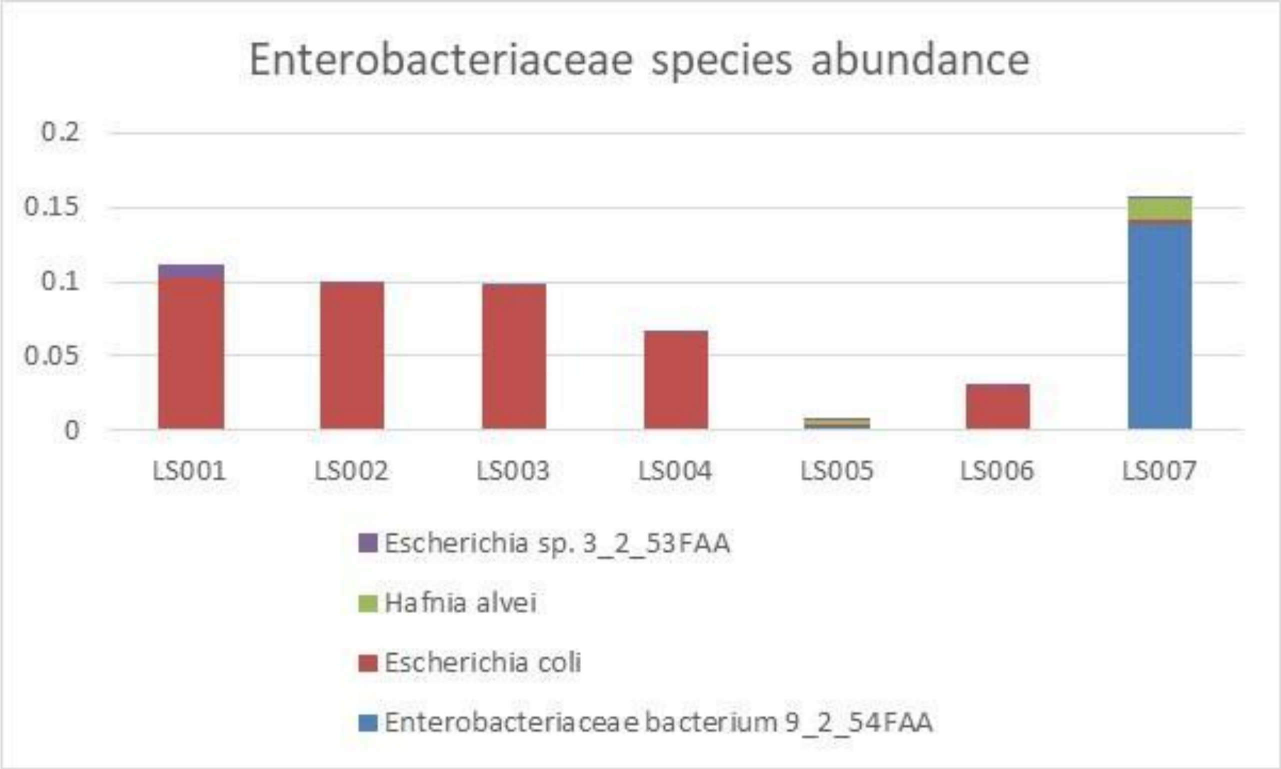


**Supplementary Figure S10:** Abundance fluctuations of the main *Enterobacteriaceae* bacteria present in the LS samples.

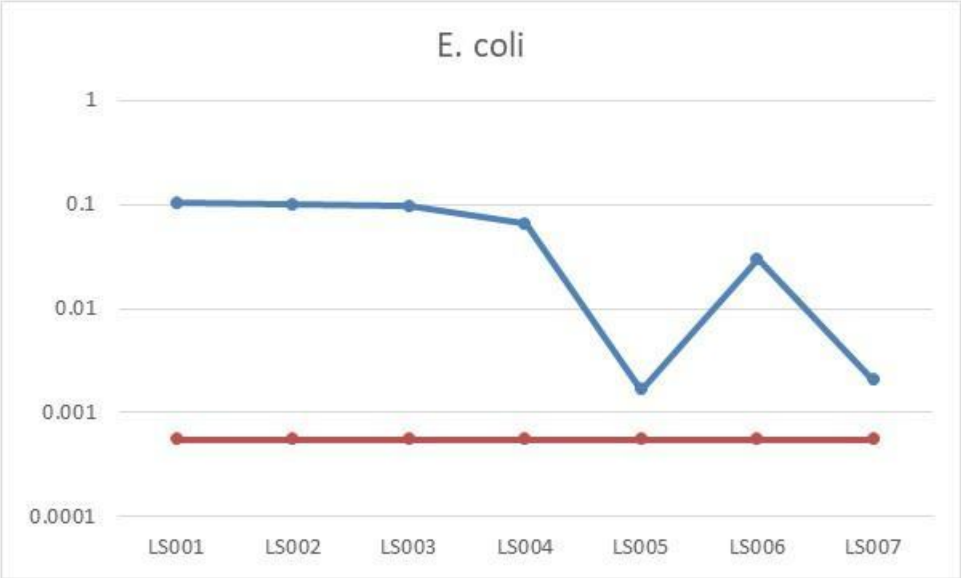


**Supplementary Figure S11:** Abundance fluctuations of E. coli in LS samples and healthy average patients.

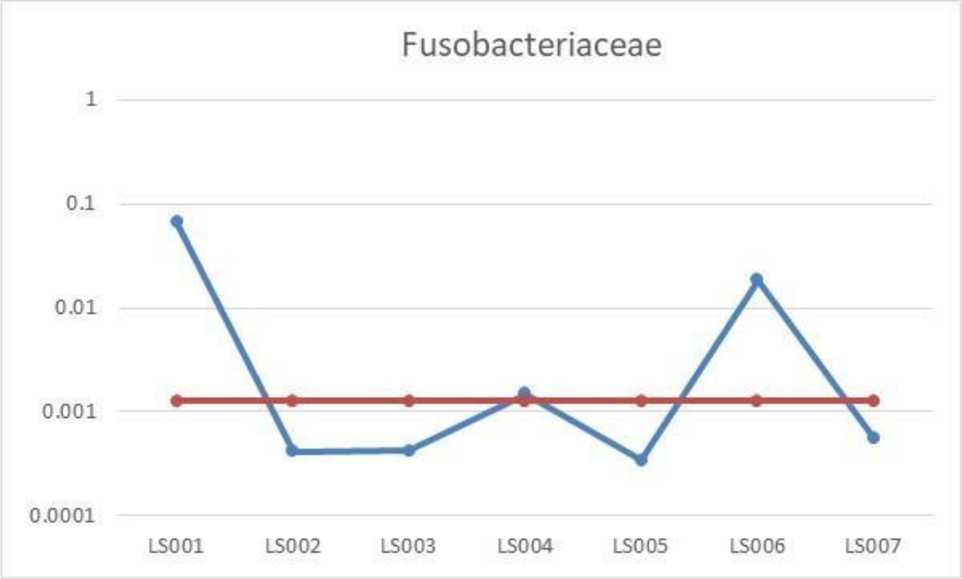


**Supplementary Figure S12:** Abundance fluctuations of microbes of the Fusobacteriaceae family in LS samples and healthy average patients.

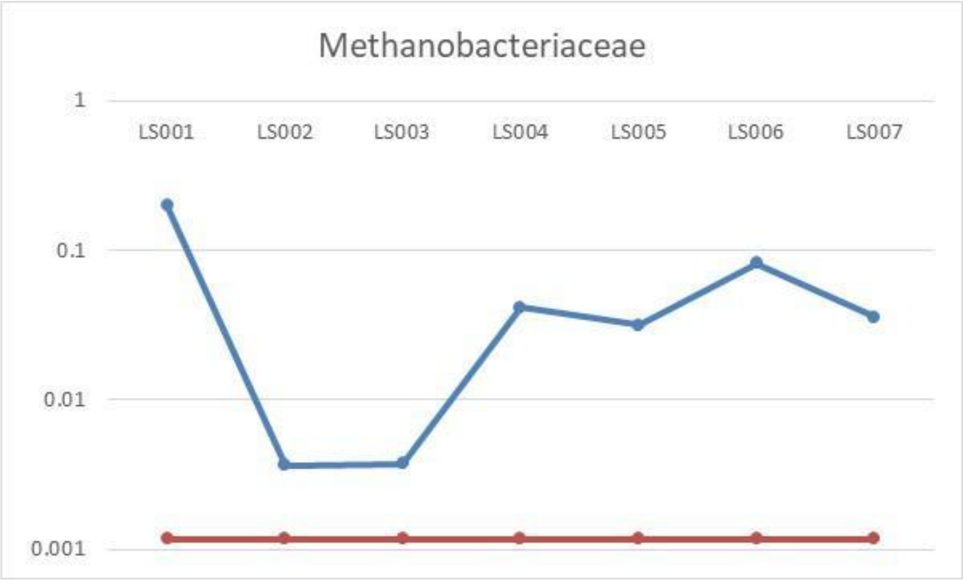

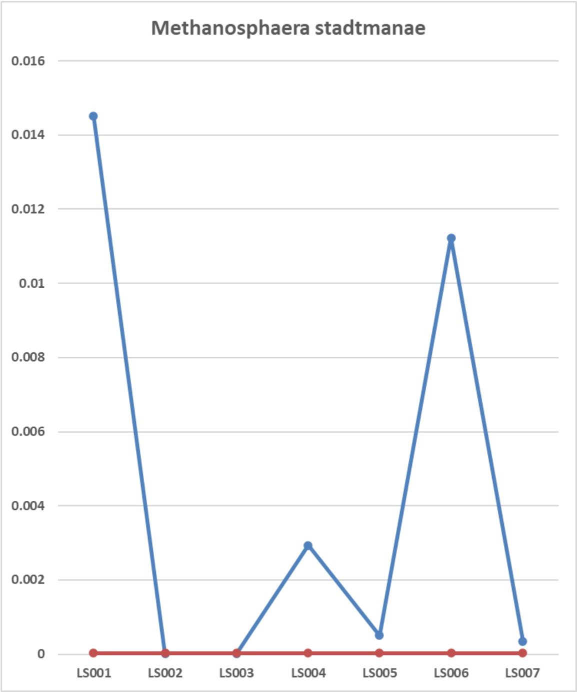


**Supplementary Figure S13:** Abundance fluctuations of microbes of the Methanobacteriaceae family in LS samples and healthy average patients, insight on *Methanosphaera sadtmanae*.

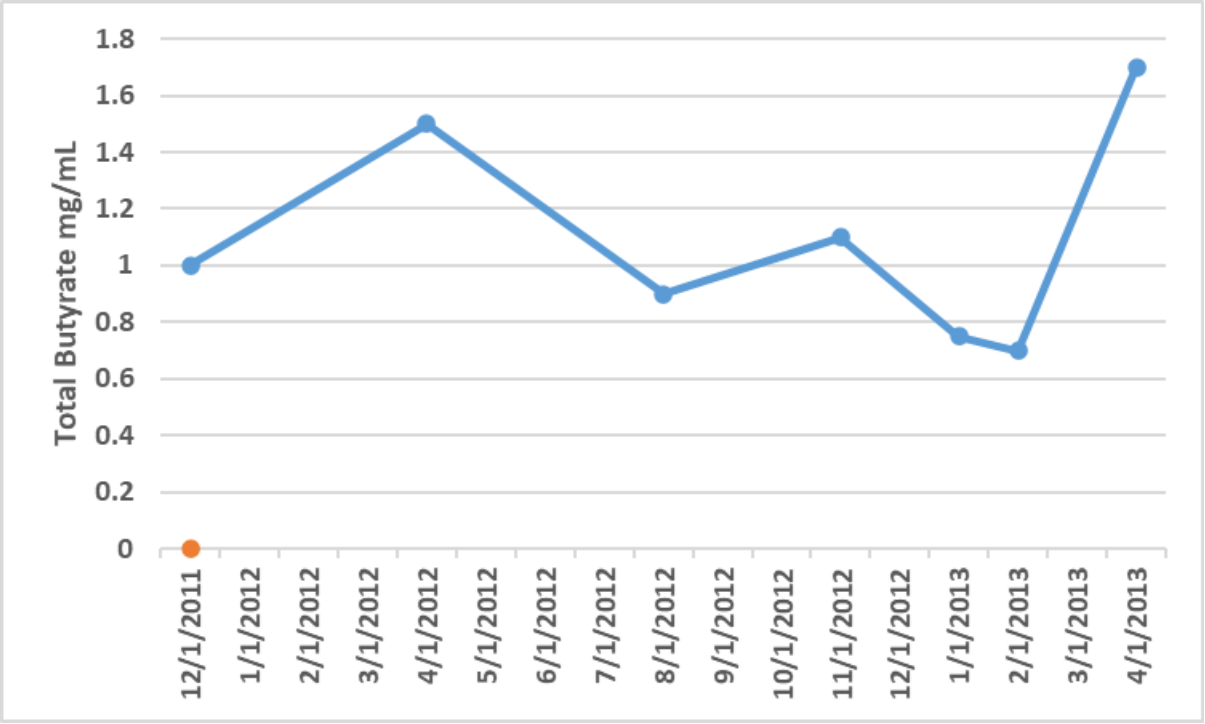


**Supplementary Figure S14:** Total butyrate from laboratory measurements on LS1-7 faecal samples.

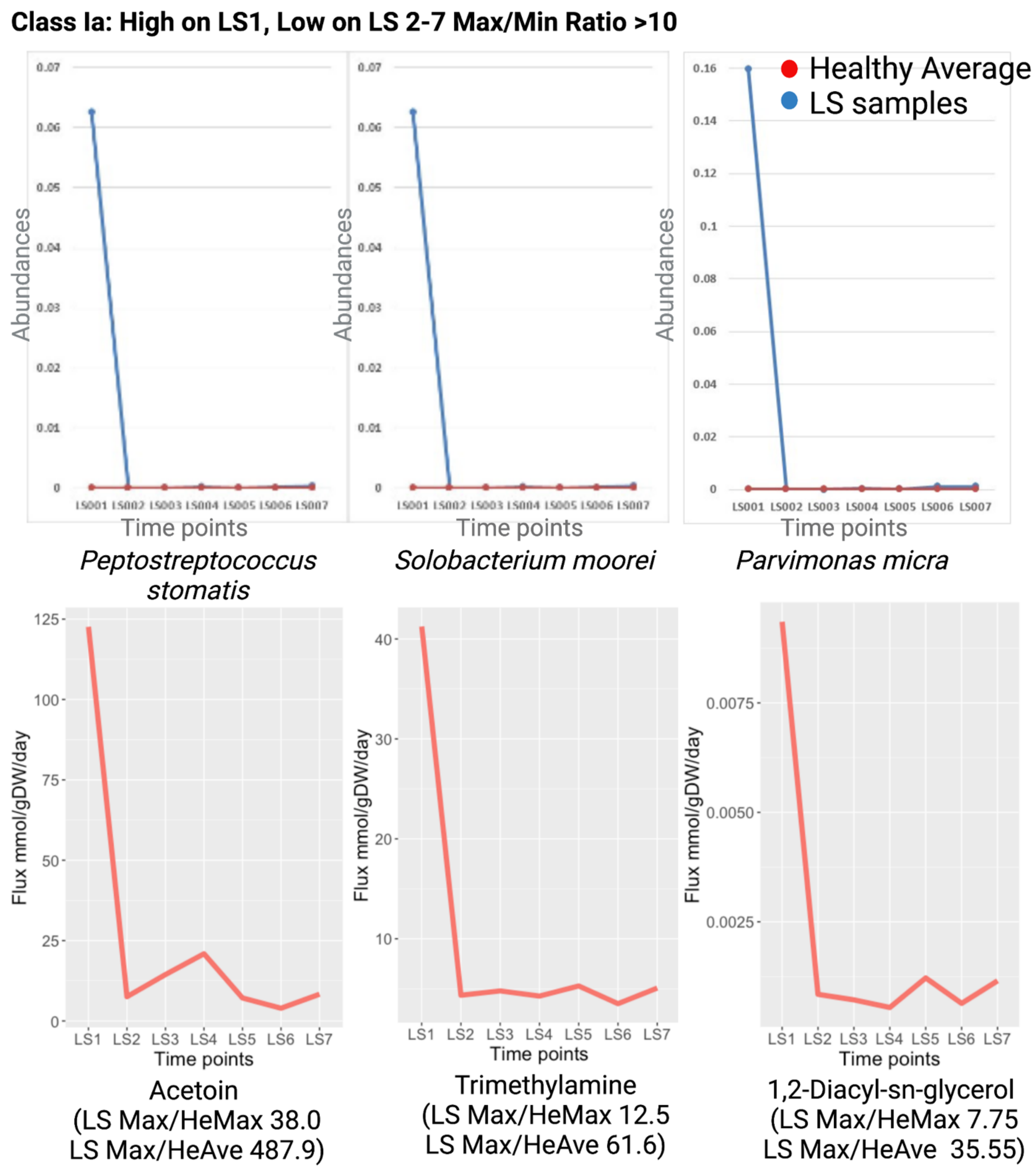


**Supplementary Figure S15:** Correlations between microbial abundances and specific fluxes, Class IA: High on LS1, Low on LS 2-7.

### Class Ib: Large Peak LS1, Low LS 2-4. Smaller Values LS5-7 Max/Min Ratio >10

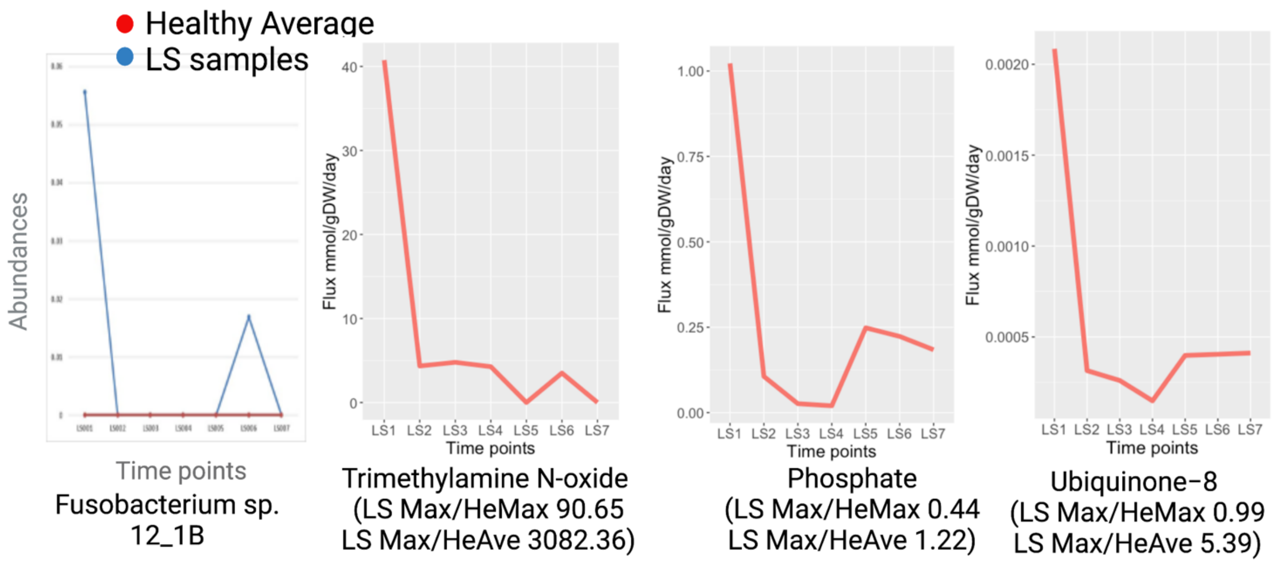


**Supplementary Figure S16:** Correlations between microbial abundances and specific fluxes, Class IB: high on LS1, other peak at LS6.

### Class Ic: High on LS1, Normal on LS 2&3, Peak Again on LS4 & LS6 Max/Min Ratio >10

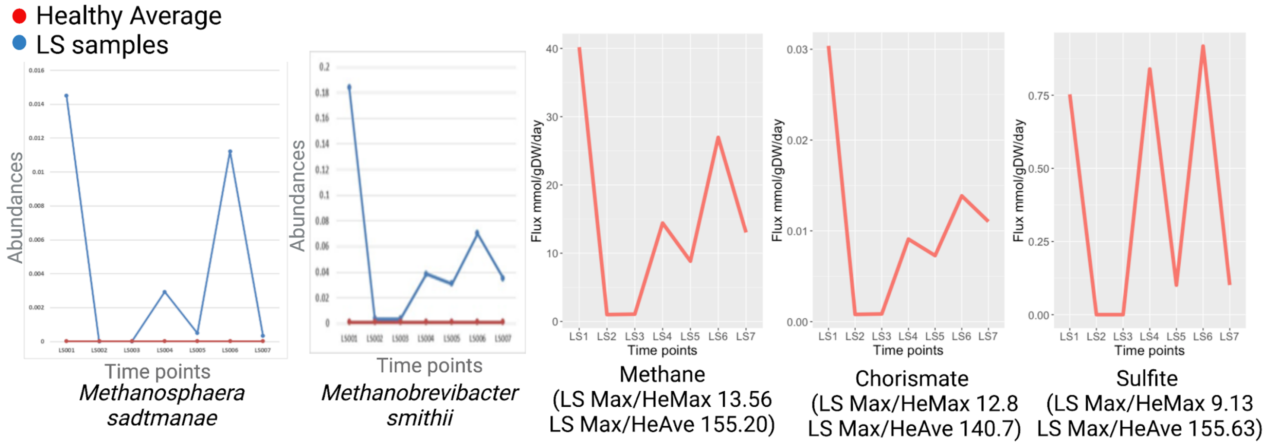


**Supplementary Figure S17:** Correlations between microbial abundances and specific fluxes, Class IC: High on LS1, normal on LS2 & 3, peak again on LS4 & LS6. Max/Min ratio >10.

### Class Id: High on LS1, 2,&3, with Another Peak at LS6, Normal on LS 5&7 Max/Min Ratio >10

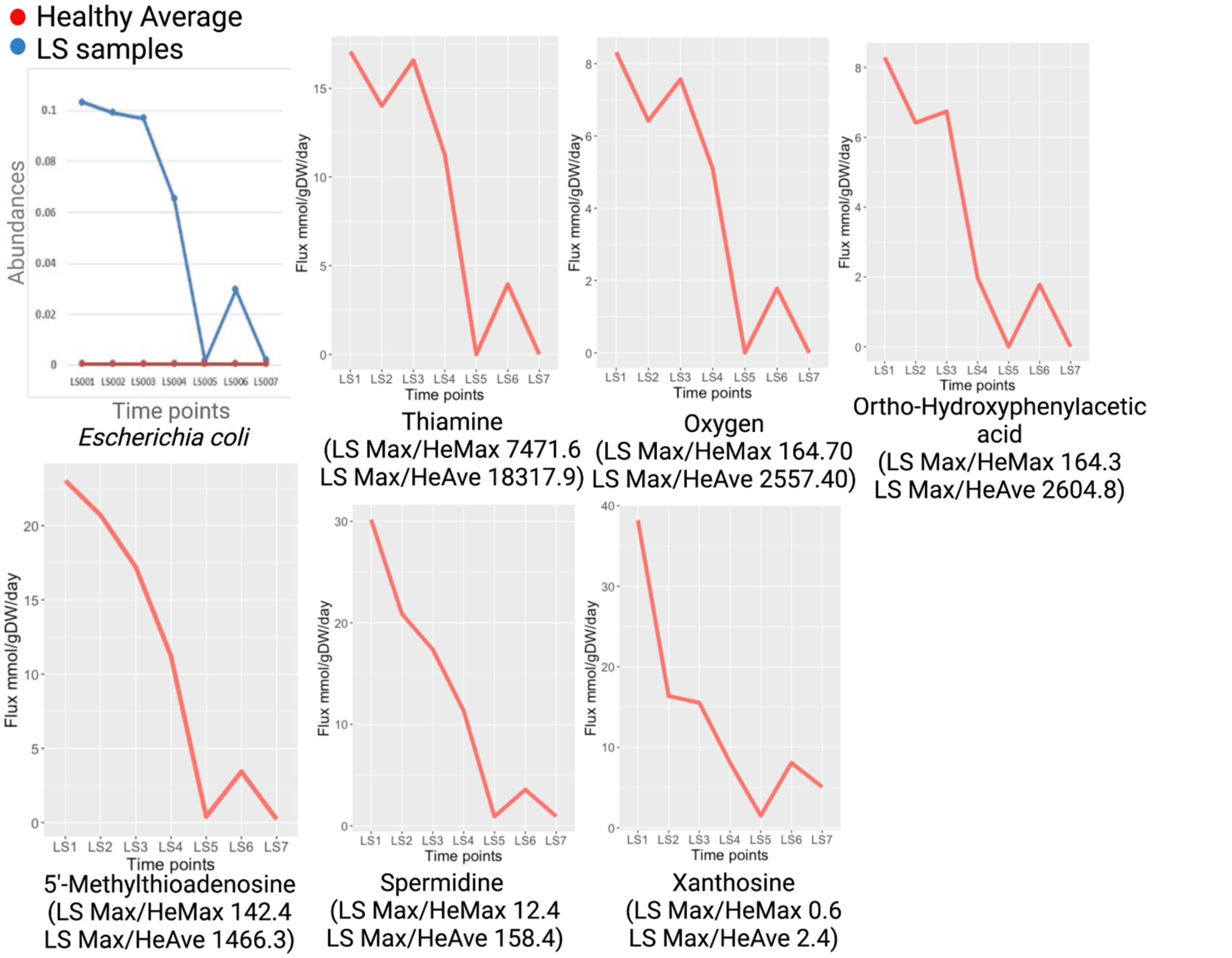


**Supplementary Figure S18:** Correlations between microbial abundances and specific fluxes, Class ID: High on LS1, 2,&3, with Another Peak at LS6, Normal on LS 5&7.

### Class Ie: High on LS1/2, LS5, LS7 Max/Min Ratio >10

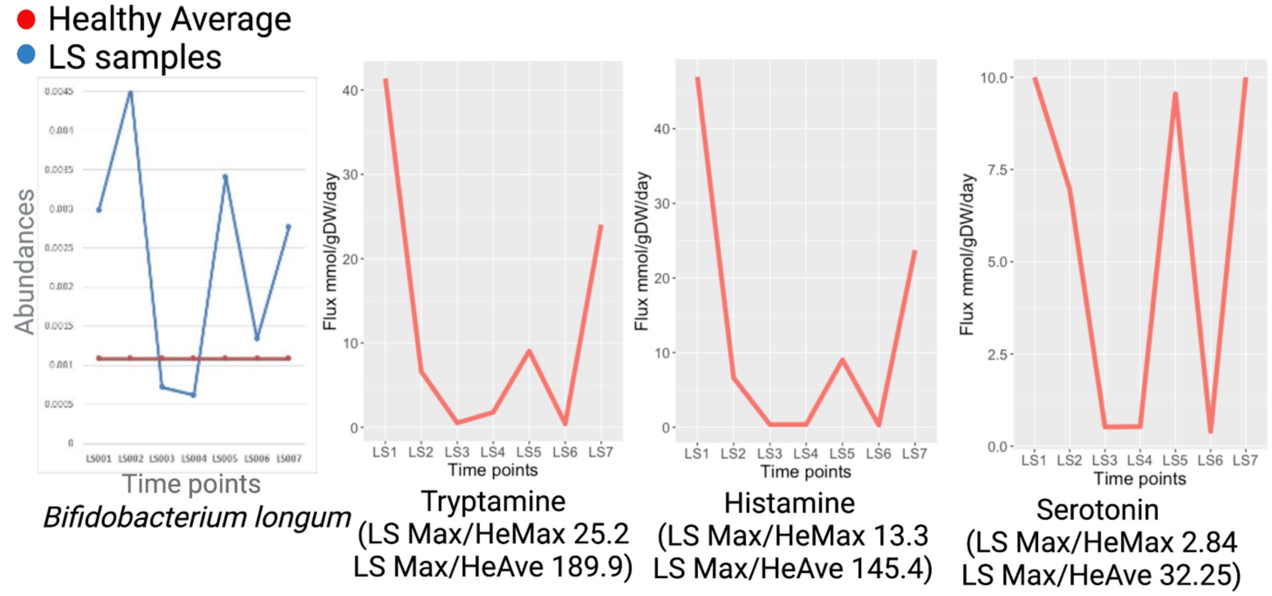


**Supplementary Figure S19:** Correlations between microbial abundances and specific fluxes, Class IE.

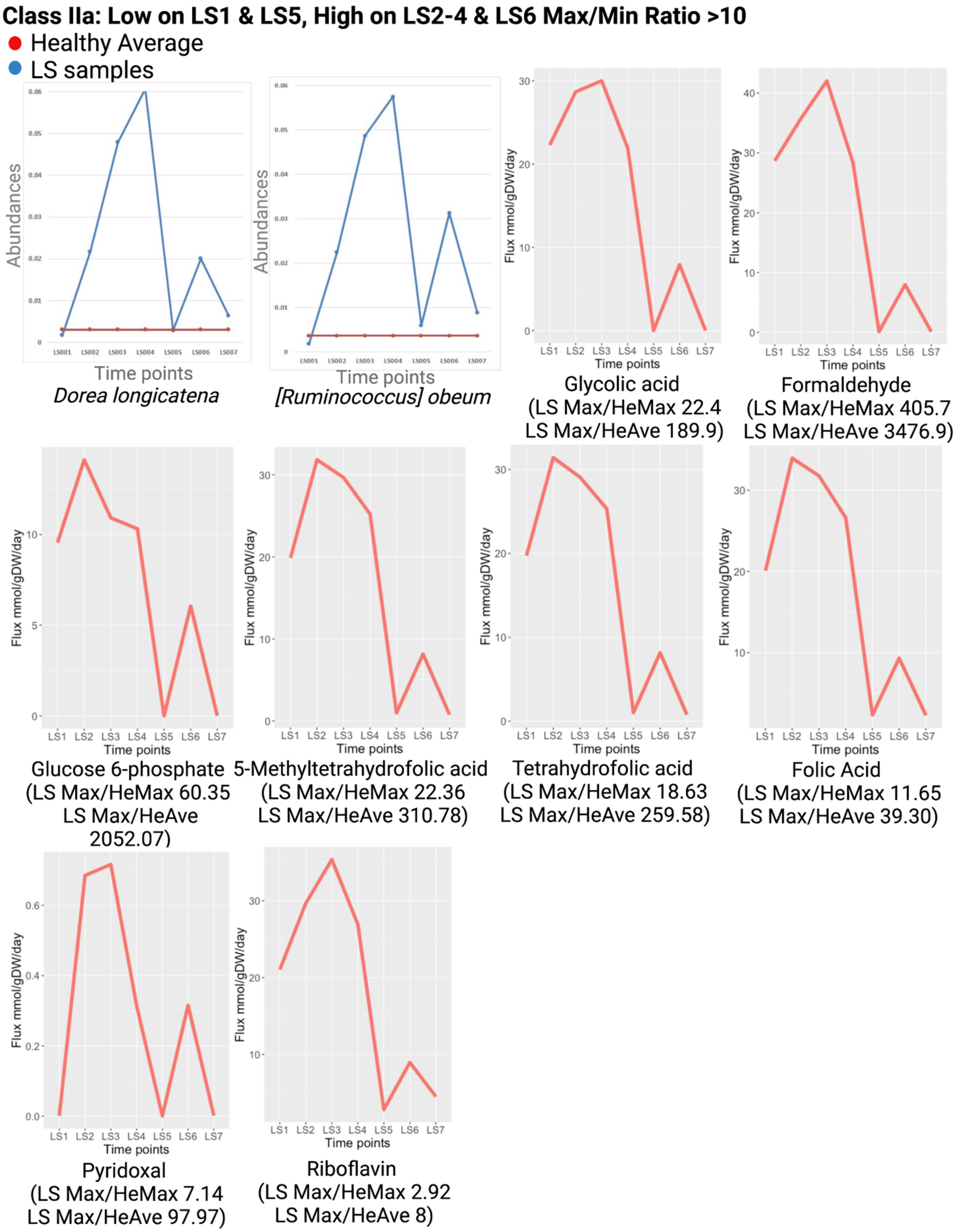


**Supplementary Figure S20:** Correlations between microbial abundances and specific fluxes, Class IIA.

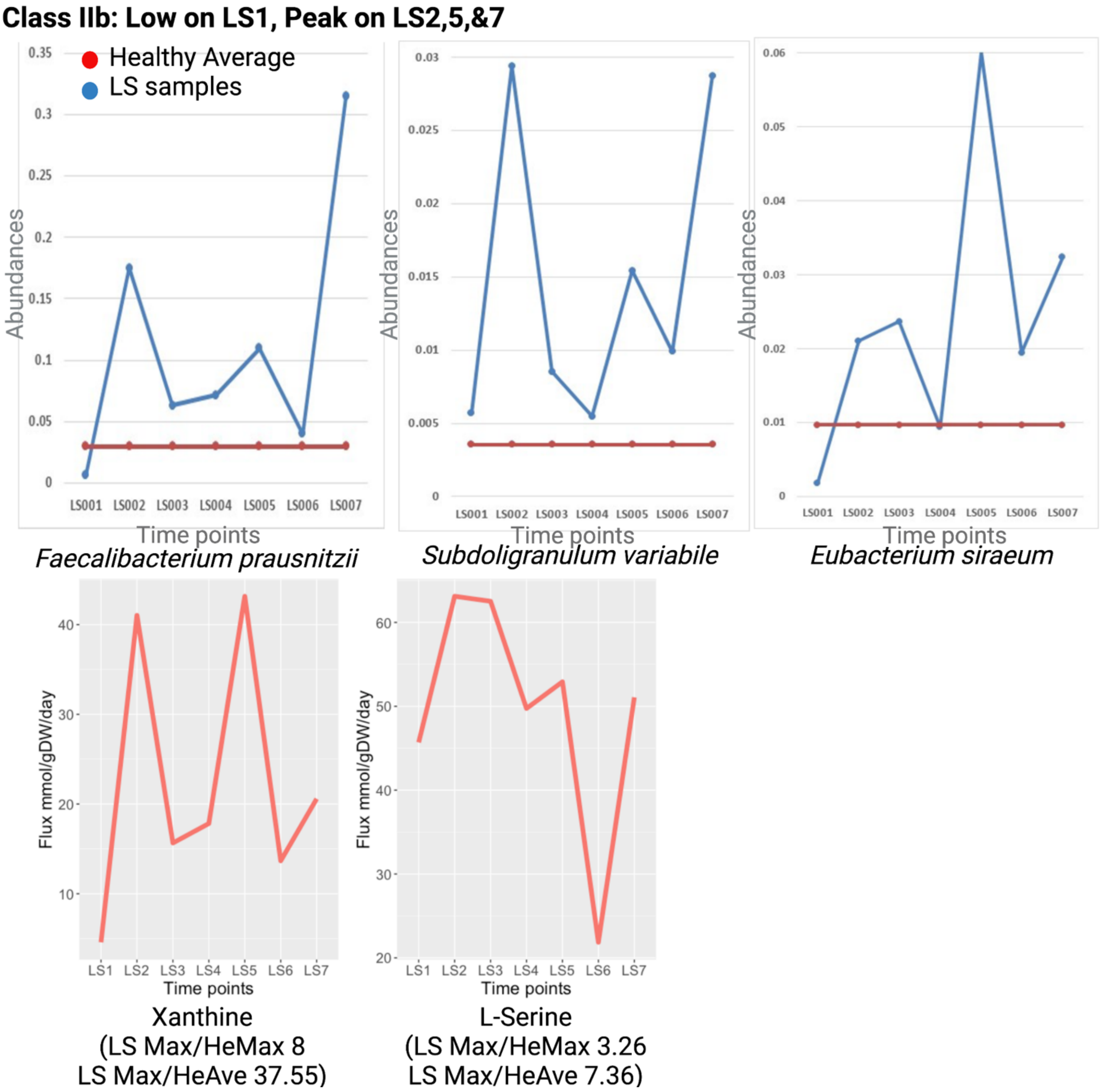


**Supplementary Figure S21:** Correlations between microbial abundances and specific fluxes, Class IIB: Low on LS1, peak on LS2-5 & LS7.
